# Synaptic Activity-Dependent Changes in the Hippocampal Palmitoylome

**DOI:** 10.1101/2021.11.26.470153

**Authors:** Glory Nasseri, Nusrat Matin, Kira Tosefsky, Greg Stacey, Stephane Flibotte, Rocio Hollman, Angela R. Wild, Leonard J. Foster, Shernaz X. Bamji

## Abstract

Dynamic protein *S*-palmitoylation is critical for neuronal function, development, and synaptic plasticity. Activity-dependent changes in palmitoylation have been observed for several neuronal substrates, however a full characterization of the activity-regulated palmitoylome is lacking. Here, we use an unbiased approach to identify differentially palmitoylated proteins in the mouse hippocampus following context-dependent fear conditioning. Of the 121 differentially palmitoylated proteins identified 63 were synaptic proteins, while others were associated with metabolic functions, cytoskeletal organization, and signal transduction. The vast majority of synaptic proteins exhibited increased palmitoylation following fear conditioning, whereas proteins that exhibited decreased palmitoylation were predominantly associated with metabolic processes. We show a link between dynamic palmitoylation and synapse plasticity by demonstrating that the palmitoylation of one of our identified proteins, PRG-1/LPPR4, is essential for activity-induced insertion of AMPA receptors into the postsynaptic membrane. Together, this study identifies networks of synaptic proteins whose dynamic palmitoylation may play a central role in learning and memory.

**SUMMARY:** This study identifies networks of proteins that undergo dynamic post-translational palmitoylation in response to fear conditioning and demonstrates that palmitoylation of one of these proteins is essential for synapse plasticity. Together, this illustrates the importance of palmitoylation in learning/memory and synapse plasticity.

## INTRODUCTION

The strengthening and weakening of synaptic connections in specific brain circuits underlies our ability to learn and remember (Takeuchi, Duszkiewicz, & Morris, 2014). Changes in synapse strength are mediated in part by dynamic post-translational modification of proteins, which in turn impact protein stability, subcellular localization and function (Citri & Malenka, 2008; Dörrbaum, Alvarez-Castelao, Nassim-Assir, Langer, & Schuman, 2020). For years, researchers have focused on activity-mediated changes in protein phosphorylation in the regulation of synapse strength (Lee, 2006; Roche, Tingley, & Huganir, 1994; Tomita, Stein, Stocker, Nicoll, & Bredt, 2005; Woolfrey & Dell’Acqua, 2015). However, it is becoming clear that other post-translational modifications, such as *S*-acylation, can play an equally important role in modulating synapse plasticity (Fukata & Fukata, 2010; Koster, 2019; Zareba-Koziol et al., 2019).

The most common form of *S*-acylation in the brain is *S*-palmitoylation (hereafter called palmitoylation), the reversible addition of the 16-carbon palmitic acid onto cysteine residue/s of a substrate protein (Chamberlain & Shipston, 2015; Ohno et al., 2012). By changing protein conformation and hydrophobicity, palmitoylation alters protein stability, protein-protein interactions, subcellular localization, and function (Aicart-Ramos, Valero, & Rodriguez-Crespo, 2011; Greaves & Chamberlain, 2007). In neurons, palmitoylation has been shown to regulate neurite outgrowth (Dumoulin, Dagane, Dittmar, & Rathjen, 2018; Lievens et al., 2016; Shah, Shimell, & Bamji, 2019), synapse formation(Shah et al., 2019; Shimell et al., 2019), synaptic transmission (Koster, 2019; Ohyama et al., 2007) and synaptic plasticity (Brigidi et al., 2014; Fukata & Fukata, 2010; Globa & Bamji, 2017; Ji & Skup, 2021; Koster, 2019). Proteomic analyses have revealed that ∼41% of all synaptic proteins are substrates for palmitoylation(Sanders et al., 2015), including scaffolding proteins (Dejanovic et al., 2014; Noritake et al., 2009; Topinka & Bredt, 1998), receptors(Van Dolah et al., 2011), ion channels (Bosmans, Milescu, & Swartz, 2011; Schmidt & Catterall, 1987; Shipston, 2013; Tian et al., 2008), cell adhesion molecules (Lievens et al., 2016), GPCRs (Kang et al., 2008; Prescott, Gorleku, Greaves, & Chamberlain, 2009; Salaun, Greaves, & Chamberlain, 2010), and proteins involved in synaptic vesicle fusion (Veit, 2000; Veit, Söllner, & Rothman, 1996). Interestingly, the dynamic palmitoylation of a subset of synaptic proteins has been reported (Brigidi, Santyr, Shimell, Jovellar, & Bamji, 2015; Fukata et al., 2013; Kaur et al., 2016; Noritake et al., 2009; Woolfrey, Sanderson, & Dell’Acqua, 2015), suggesting that this post-translational modification may be important for synapse plasticity. Indeed, PSD-95(Noritake et al., 2009), δ-catenin (Brigidi et al., 2015), and A-kinase anchoring protein 79/150 (Woolfrey et al., 2015) are dynamically palmitoylated in response to changes in synaptic activity, and palmitoylation of each has been shown to be critical for the recruitment of glutamatergic AMPA receptors (AMPARs) to the postsynaptic membrane, the enlargement of dendritic spines, and activity-induced synapse strengthening (Bredt & Nicoll, 2003; Matsuzaki, Honkura, Ellis-Davies, & Kasai, 2004).

Dysregulated palmitoylation has been implicated in the pathophysiology of a variety of neurological and psychiatric diseases, including schizophrenia, intellectual disability and Alzheimer’s disease (Cho & Park, 2016; Sanders et al., 2015; Zareba-Koziol et al., 2019). Therefore, understanding how aberrations in protein palmitoylation affect synaptic plasticity could facilitate a better understanding of the etiology of these developmental disorders. While palmitoylation acts as a critical molecular regulator of intracellular protein dynamics and function in the brain, it remains unclear: 1) which substrates are differentially palmitoylated in response to fluctuations in synaptic activity; and 2) whether the differential palmitoylation of these proteins is important for activity-induced changes in synapse strength.

In the present study, we use a proteomic approach to identify proteins that are differentially palmitoylated after a learning event. We generated a list of 121 proteins whose palmitoylation levels were either significantly altered in the fear conditioned group compared to control or showed a greater than 2-fold difference between these two conditions. Following the validation of a number of proteins from the proteomic screen, we focused on a single protein identified in our list, PRG-1/LPRR4 (plasticity-related gene 1/ lipid phosphate phosphatase-related protein type 4). PRG-1 is a postsynaptic protein found predominantly in glutamatergic neurons that modulates the release probability of presynaptic glutamate vesicles at excitatory synapses (Liu et al., 2016; Tokumitsu et al., 2010; Trimbuch et al., 2009) and has been implicated in regulating hippocampal synaptic plasticity and spatial memory formation(Liu et al., 2016). We demonstrate that palmitoylation of the canonical 83 kDa isoform of PRG-1 is essential for spine formation and for activity-induced insertion of AMPA receptors into the postsynaptic membrane. Together, this study identifies novel networks of proteins that are palmitoylated in an activity-dependent manner following a learning event and suggests a key role for the differential palmitoylation of synaptic proteins in synapse plasticity and memory acquisition.

## RESULTS

### Identification of differentially palmitoylated hippocampal proteins

A number of *in vitro* and *in vivo* studies have previously described the differential palmitoylation of proteins in response to changes in synaptic activity (Brigidi et al., 2015; Dejanovic et al., 2014; Hanley & Henley, 2010; Hayashi, Rumbaugh, & Huganir, 2005; Kaur et al., 2016; Thomas, Hayashi, Chiu, Chen, & Huganir, 2012; Van Dolah et al., 2011; Woolfrey et al., 2015). While it is clear that the palmitoylation of synaptic proteins is important in regulating protein function, a more complete picture of *which* proteins are dynamically palmitoylated and *how* the dynamic palmitoylation of proteins contributes to synaptic plasticity is lacking. We therefore took an unbiased, proteomic approach to identify proteins that are differentially palmitoylated in response to contextual fear-conditioning (FC). We first confirmed acquisition of contextual memory by demonstrating increased freezing behavior in 9-week-old male mice 1 hour after a single foot shock (0.3mA, 5 sec) (Fig. 1 A). Mouse hippocampi were then immediately collected, the membrane fraction enriched, and palmitoylated proteins isolated using the acyl-resin assisted capture (acyl-RAC) assay (Fig. 1, B and C). In this assay, free cysteine residues were first blocked by incubating lysates with methyl methanethiosulfonate (MMTS). Next, the palmitoyl-thioester bond was cleaved using hydroxylamine (HAM), resulting in the exposure of a free sulfhydryl that can bind to the sulfhydryl-reactive Sepharose resin (Fig. 1 B)(Forrester et al., 2011). To control for non-specific binding of proteins to the resin, the samples were divided into equal parts and half the sample was processed without the HAM cleavage step [(-)HAM] (Fig. 1 C). For each control and FC animal, 3 fractions were analyzed by label-free quantification (LFQ) liquid chromatography-mass spectrometry (LC-MS/MS): i) input (lysate prior to acyl-RAC assay); ii) the (+)HAM fraction, which represents palmitoylated proteins captured by the resin; and iii) the (-)HAM fraction, which identifies proteins that bound non-specifically to the resin (Fig. 1 C).

**Figure 1:**
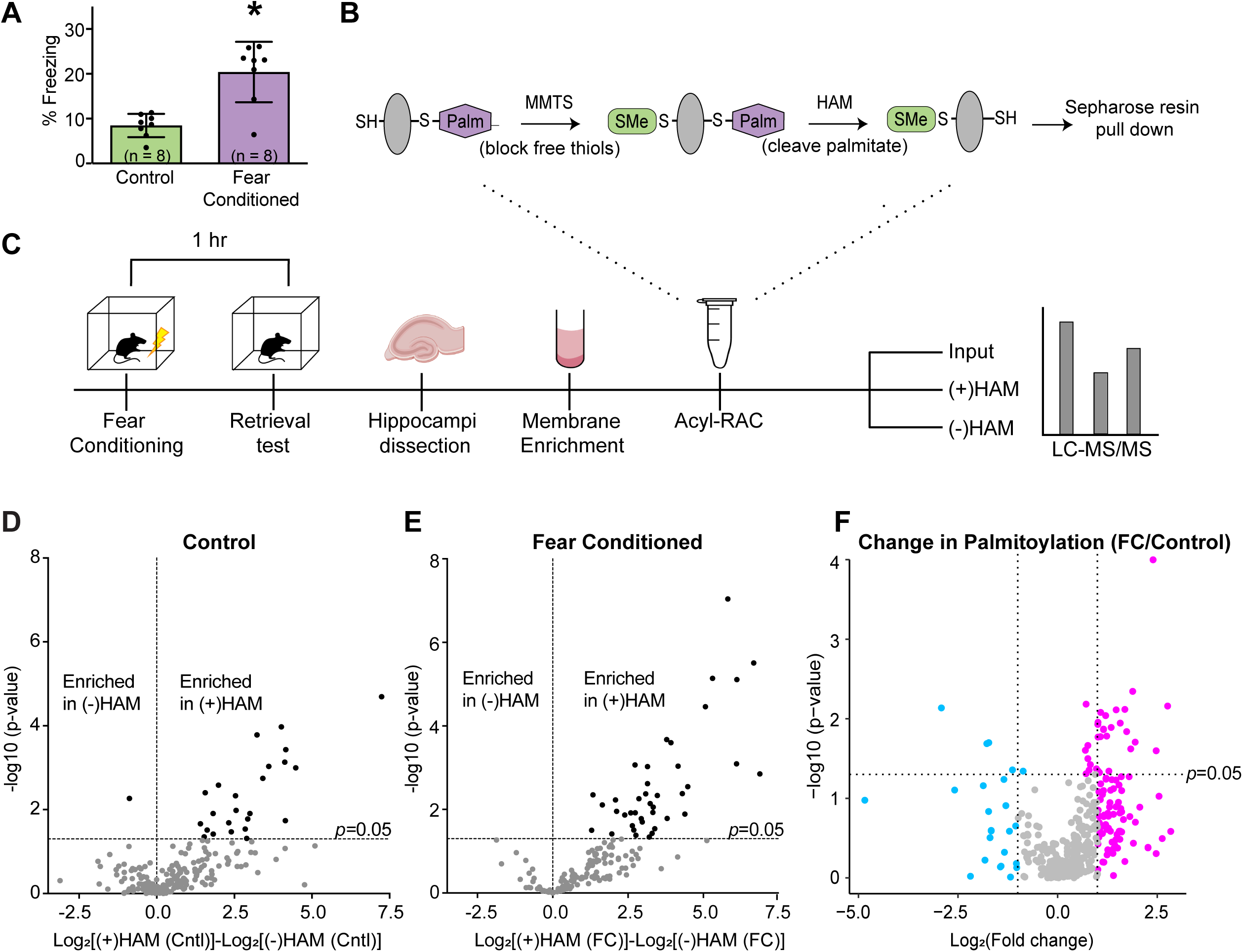
Contextual fear conditioning results in the differential palmitoylation of 121 proteins. **(A)** Fear-conditioned mice exhibited greater freezing behavior than sham control, demonstrating efficacy of the fear conditioning. **(B)** Schematic representation of the acyl-RAC assay used to isolate palmitoylated proteins. **(C)** Schematic illustration of the experimental design: a memory retrieval test was done 1 hr following fear conditioning and hippocampi immediately collected. Membrane-enriched samples were run through the acyl-RAC assay to isolate palmitoylated proteins. Inputs, (+)HAM, and (-)HAM samples were run on LFQ LC-MS/MS to identify and quantify palmitoylated proteins. FC=fear conditioning; HAM=hydroxylamine; LC-MS/MS = liquid chromatography-tandem mass spectrometry. **(D and E)** Overrepresentation of proteins in the (+)HAM control **(D)** and (+)HAM FC **(E)** samples, relative to their (-)HAM samples. Graphs depict fold-change in protein abundance (x-axis) and the significance of the enrichment in the (+)HAM sample compared to (-)HAM (y-axis, two-sided t-test). **(F)** Fold change in the abundance of palmitoylated proteins in FC samples compared to control. Each dot represents an individual protein. Pink dots represent proteins that exhibit increased palmitoylation (either >2-fold increase - right of vertical line, or significantly (p < 0.05, two-sided t-test) different - above horizontal line) and blue dots represent proteins that exhibit decreased palmitoylation (> 2-fold decrease - left of vertical line, or significantly (p < 0.05, two-sided t-test) different - above horizontal line) in the FC group compared to control.

In total, we identified 1940 proteins throughout all experimental conditions (Table S1). After additional filtering, 345 proteins with at least 2 LFQ values in control and FC (+)HAM groups were analyzed (Table S2). Since the missing values were randomly distributed across the samples, pairwise correlation analyses were done to assess reproducibility (Perrin et al., 2013), with representative graphs from control and FC groups shown in Figures S1 A-C. Our results demonstrate strong correlation between biological replicates with respect to LFQ intensities (Fig. S1 A; average R= 0.73) and thus that the sequential nature of mass spectroscopy did not systematically bias protein quantification in our samples.

To ensure that any detected changes in protein palmitoylation were not attributable to changes in protein turnover, we compared the input fractions from fear-conditioned and sham mice. Of the 1163 proteins with at least two quantified LFQ intensities in both input fractions, only 45 (4%) showed a significant change in input fraction levels between the fear-conditioned and sham group (p < 0.05, two-sided t-test). No *p*-values passed multiple comparison correction (Benjamini-Hochberg), indicating that there were no systematic differences in protein input levels between the two groups.

We next compared enrichment in the (+)HAM compared to the (-)HAM fractions as a further test for false-positives (i.e. proteins that bound non-specifically to the resin) (Fig. 1, D and E), as well as (+)HAM control vs (+)HAM FC to identify proteins that were differentially palmitoylated following FC (Fig. 1 F). While a fraction of proteins did appear to be enriched in both control and FC (-)HAM fractions (Fig. 1 D and E; points left of the dotted line), only one protein in the control sample and no proteins in the FC sample were significantly enriched in the (-)HAM fraction (left of the dotted line and above the p=0.05 line). This demonstrates a low rate of false positives in our assay. Of the 345 palmitoylated proteins quantified in our screen, we focused on 121 proteins of interest: 35 proteins whose palmitoylation levels differed significantly (p < 0.05) between control and FC groups (Fig. 1 F, above the p=0.05 line), and an additional 86 proteins that exhibited a greater than two-fold change in palmitoylation levels between the two groups (Fig. 1 F, outer left and right of the vertical dotted lines) (Table S3). 72.5% of proteins in the FC sample exhibited an increase (Fig. 1 F, pink dots) and 27.5 % exhibited a decrease (Fig. 1 F, blue dots) in palmitoylation 1 hour after FC.

The 121 proteins of interest were grouped by their main parent term categories based on either their associated ‘biological process’ or ‘molecular function’ gene ontology (GO) annotation terms using the mouse genome informatics (MGI) database (version 6.15; http://www.informatics.jax.org) (Bult, Blake, Smith, Kadin, & Richardson, 2019) (Table S3).

These proteins fell into 22 classifications including many processes critical to the regulation of synaptic plasticity, such as “synaptic vesicle cycle” (GO:0099504), “cell adhesion” (GO:0007155), “cytoskeleton organization” (GO:0007010), “receptor clustering” (GO:0043113), “GTPase activity” (GO:0003924) and “ion transport” (GO:0006811).

To determine which synaptic proteins are subject to activity-induced palmitate turnover, we analyzed our list of 121 differentially palmitoylated proteins using the synaptic protein database, SynGO (Koopmans et al., 2019). We found that 63 of our 121 differentially palmitoylated input genes mapped to one of the 1,225 SynGO annotated genes found within a “brain expressed” background set of 18,035 unique genes. Both the SynGO (Koopmans et al., 2019) and MGI (Mouse Genome Informatics) (Bult et al., 2019) databases were used to illustrate the subcellular localization of the identified proteins at the synapse (Fig. S2). An additional 7 astrocyte-enriched proteins from our list were identified in the Astrocyte Transcriptome database (Cahoy et al., 2008) and highlighted in Figure S2.

We examined the relationships between our 121 proteins of interest using STRING protein-protein interaction network analysis (Fig. 2 A) (version 11.0, released 2017-05-14) (Szklarczyk et al., 2019; von Mering et al., 2005). STRING analysis, with parameters set to a medium confidence level of 0.40, revealed 413 edges representing direct and indirect interactions between these 121 proteins, with an enrichment *p*-value of < 1 x 10 ^-16^. Within the STRING network, we noticed two distinct groups: the first comprising 21 proteins that mapped to the “metabolic process” (GO:0008152) parent GO term (Fig. 2 B; Table S3) and the second containing 63 proteins which mapped to synaptic cellular component and/or biological process terms in SynGO (Koopmans et al., 2019) (Fig. 2 C). 83% of synaptic proteins exhibited increased palmitoylation following FC, while conversely, most metabolic proteins (57%) exhibited decreased palmitoylation following FC. Majority of brain metabolic energy is used on synaptic transmission and changes in synapse strength can alter energy expenditure (Harris, Jolivet, & Attwell, 2012). As mitochondrial enzymes have been shown to be inhibited by palmitoylation (Berthiaume, Deichaite, Peseckis, & Resh, 1994; Corvi, Soltys, & Berthiaume, 2001), the activity-induced *reduction* in metabolic protein palmitoylation observed in this study may serve a regulatory function to *enhance* enzymatic activity during neuronal stimulation and synaptic transmission.

**Figure 2:**
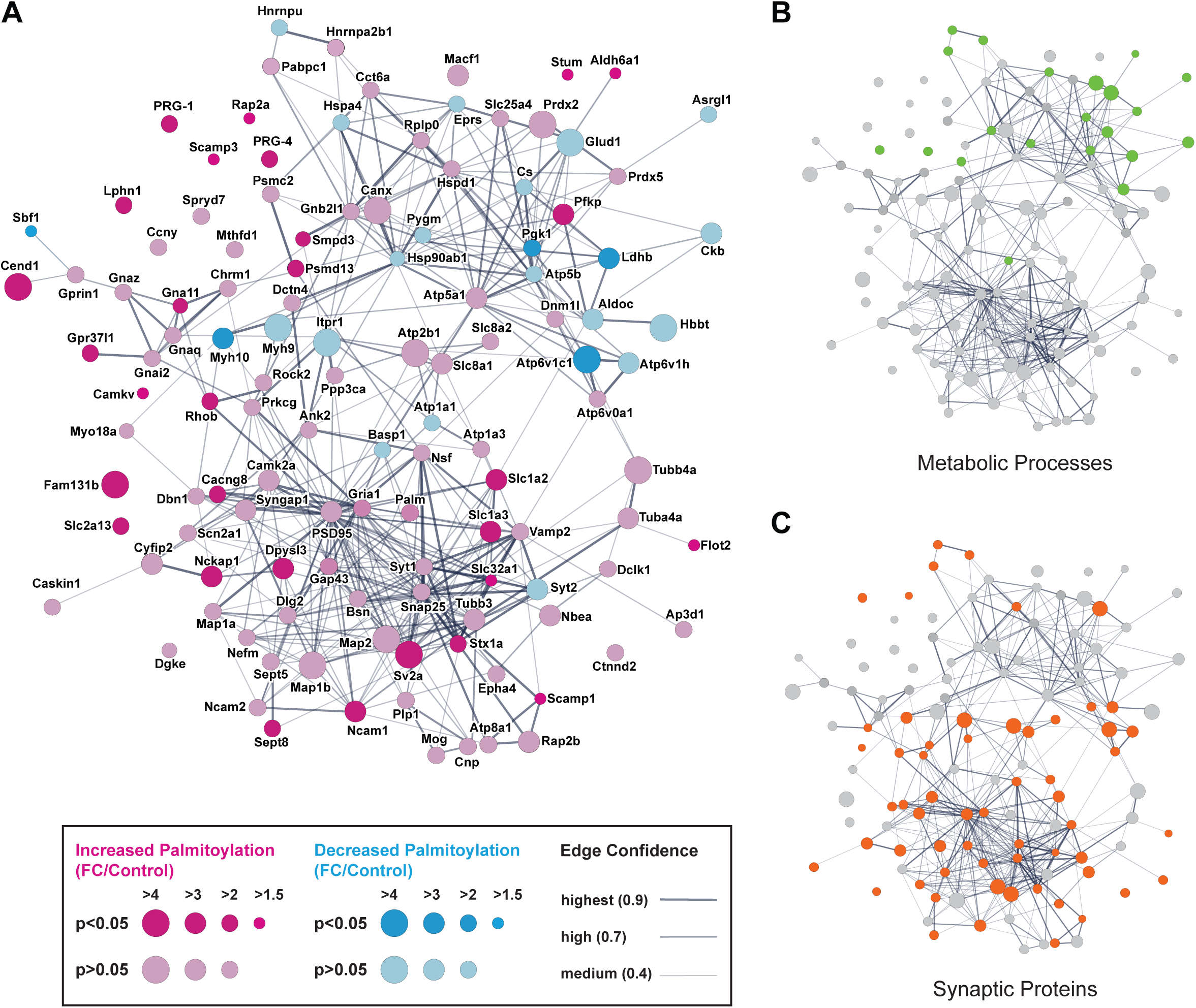
STRING protein-protein interaction network analysis. **(A)** STRING analysis of the 121 proteins differentially palmitoylated after FC. Each node represents all the proteins produced by a single, protein-coding gene locus and the number of edges represent protein-protein associations. Thickness of the edges indicate strength of data support. STRING parameters were set to medium confidence (0.400) and analyzed against the mouse genome. **(B)** 21 proteins which mapped to metabolic process gene ontology terms in MGI(Bult et al., 2019) are highlighted in green. Of these metabolic proteins, 12 exhibited decreased palmitoylation levels. **(C)** 63 proteins mapped to synaptic biological process or cellular component gene ontology terms in SynGO(Koopmans et al., 2019), majority of which (52 of 63) showed an increase in palmitoylation, and are highlighted in orange.

Consistent with our GO analysis, Kyoto Encyclopedia of Genes and Genomes (KEGG) (Kanehisa & Sato, 2020) analysis (Fig. S3; Table S4) revealed “pathways of neurodegeneration” (19 genes), “metabolic pathways” (19 genes), “synaptic vesicle cycle” (11 genes),“calcium signaling pathway” (11 genes) and “glutamatergic synapse” (9 genes) as among the terms most overrepresented in our gene list. Similar disease associations were found via disease enrichment analysis using DisGeNET v7 (Piñero et al., 2019), where “Schizophrenia”, “Bipolar Disorder”, “Alzheimer’s disease” and “Major Depressive Disorder” emerged as the diseases most strongly associated with our list of differentially palmitoylated proteins (Fig. S4; Table S5).

### Co-expression of differentially palmitoylated proteins with known palmitoylation enzymes

To investigate specific enzymes that may be responsible for mediating activity-dependent palmitate turnover, we examined the co-expression of our 121 substrate proteins of interest with the known ZDHHC enzymes, accessory proteins, and the best characterized de-palmitoylating enzymes (30 genes total). We extracted single-cell RNAseq expression data for the mouse hippocampus from DropViz (Saunders et al., 2018), performed Spearman correlation analysis and plotted a co-expression matrix with hierarchical clustering (Fig. 3 A). Our analysis revealed numerous significant correlations (Table S6 for statistical measures, FDR < 0.05), most notably a subset of 10 enzymes and accessory proteins (*Zdhhc2, Zdhhc3, Zdhhc5, Zdhhc8, Zdhhc13, Zdhhc17, Zdhhc21, Zdhhc23*, *Golga7b* and *Lypla2*) that co-expressed highly with 73 of the 121 differentially palmitoylated substrates (Fig 3 A, dotted box).

**Figure 3:**
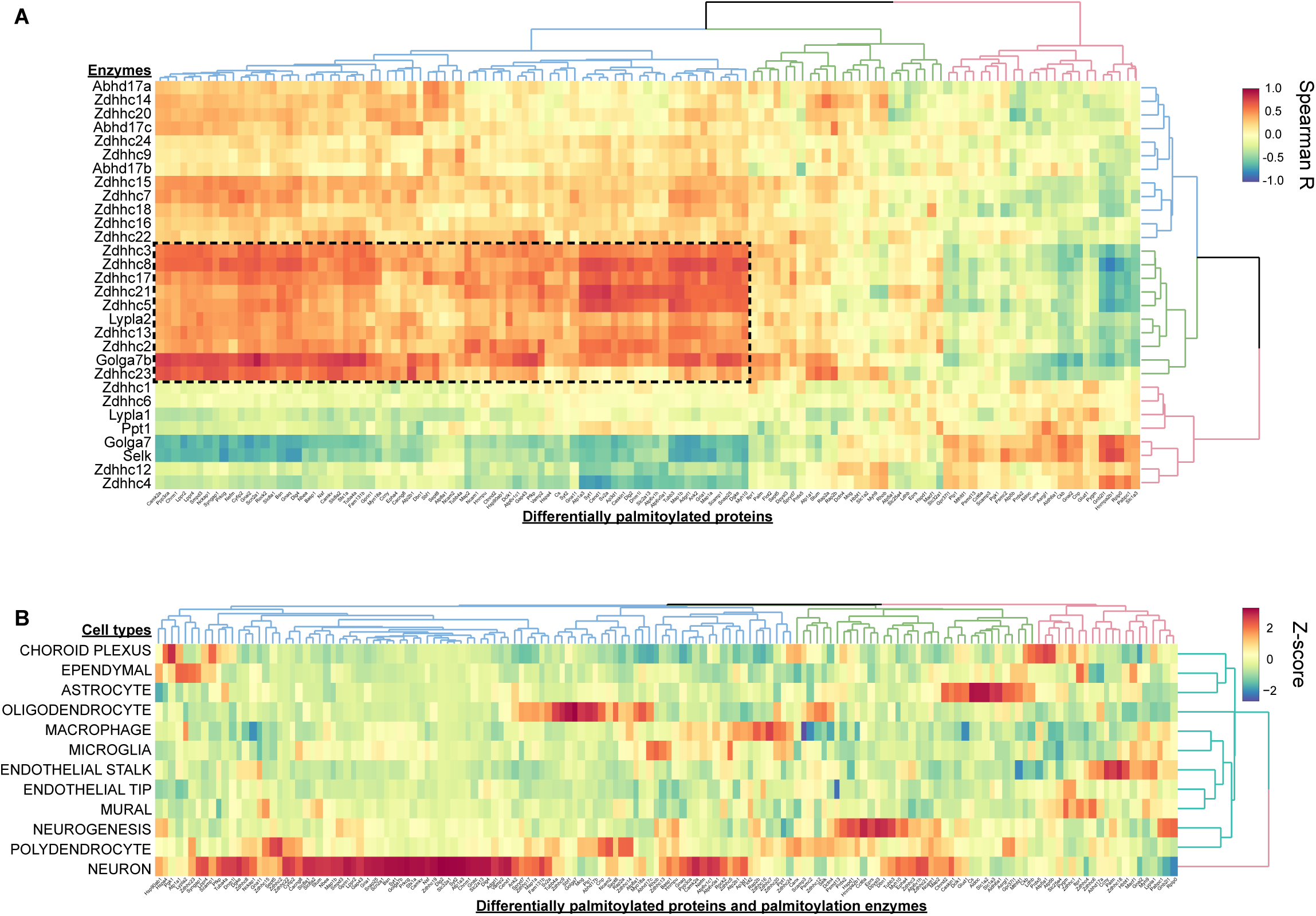
Co-expression analyses of palmitoylating / depalmitoylating enzymes and the 121 differentially palmitoylated proteins. **(A)** Heatmap depicting the co-expression of the 121 differentially palmitoylated substrates with 30 palmitoylating/depalmitoylating enzymes and accessory proteins. Spearman correlation values between genes were calculated using expression data from DropViz across 103 hippocampal cell classes. 10 enzymes and accessory proteins were highly co-expressed with 73 differentially palmitoylated substrates (dotted box comprising the blue branch of the x-axis dendrogram and the green branch of the y-axis dendrogram). **(B)** Heatmap depicting the expression of the 30 enzymes/accessory proteins and 121 differentially palmitoylated substrates in different hippocampal cell types. Z scores were calculated using mean expression values from DropViz for each cell class.

We also determined whether the highly correlating enzymes/accessory proteins and substrates (Fig. 3 A, dotted box) were specifically enriched in neurons. We generated a heatmap of average cell-type expression z-scores from DropViz (Saunders et al., 2018) for the set of 30 palmitoylation associated genes and 121 substrates and found that the majority of differentially palmitoylated substrates were highly enriched in neurons (71 of 121 genes, Fig. 3 B, Table S7; neuronal z-score > 1). Notably, 49 of the 83 highly co-expressed genes in Fig 1 A (10 enzymes/accessory proteins and 73 substrates, dotted box) were enriched in neurons (Fig. 3 B). These results indicate that this subset of enzymes/accessory proteins may act as the predominant palmitoylation machinery in hippocampal neurons and mediate the palmitoylation of neuron enriched substrates identified in our study.

In addition, we observed several clusters of activity-regulated palmitoylation substrates that were highly enriched in a variety of non-neuronal cell types, including oligodendrocytes, astrocytes, and endothelial cells, indicating that differential palmitoylation may also be important for plasticity-related signaling in other cell types. The raw correlation matrix and expression matrix can be found in Tables S6 and S7, respectively.

### Validation of differentially palmitoylated hippocampal proteins

To validate that the 121 proteins identified in our proteomic screen following FC are differentially palmitoylated in an activity-induced manner, we tested whether chemical long-term potentiation (cLTP) resulted in the differential palmitoylation of a subset of these proteins in primary hippocampal neuron cultures. Our lab has previously demonstrated an increase in the palmitoylation of δ-catenin 1 hour following cLTP *in vitro* which was recapitulated 1 hour following fear conditioning *in vivo* (Brigidi et al., 2014). Primary hippocampal cultures were used instead of whole hippocampi in order to focus on differentially palmitoylated proteins that are more directly involved in synapse strengthening in neurons. Specifically, 14 days *in vitro* (DIV) cultures were treated with a standard glycine cLTP protocol shown to enhance synaptic activity and spine size through the selective activation of synaptic NMDA receptors (NMDARs) (Lu et al., 2001). One hour later, lysates were collected, enriched for the membrane fraction, and palmitoylated proteins isolated using the acyl-RAC assay (Fig. 4 A). We focused on 13 of the 121 proteins based on the robustness of the differential palmitoylation following FC and the quality of available antibodies. 8 of the 13 tested proteins exhibited differential palmitoylation in response to cLTP as well as FC (Fig. 4 B), whereas 5 of the 13 candidates did not (Fig. 4 C). These results indicate that dynamic changes in protein palmitoylation are reliably induced following a variety of synaptic stimulation protocols and are therefore likely to play an important role in coordinating common signaling events downstream of different modes of synaptic activation.

**Figure 4:**
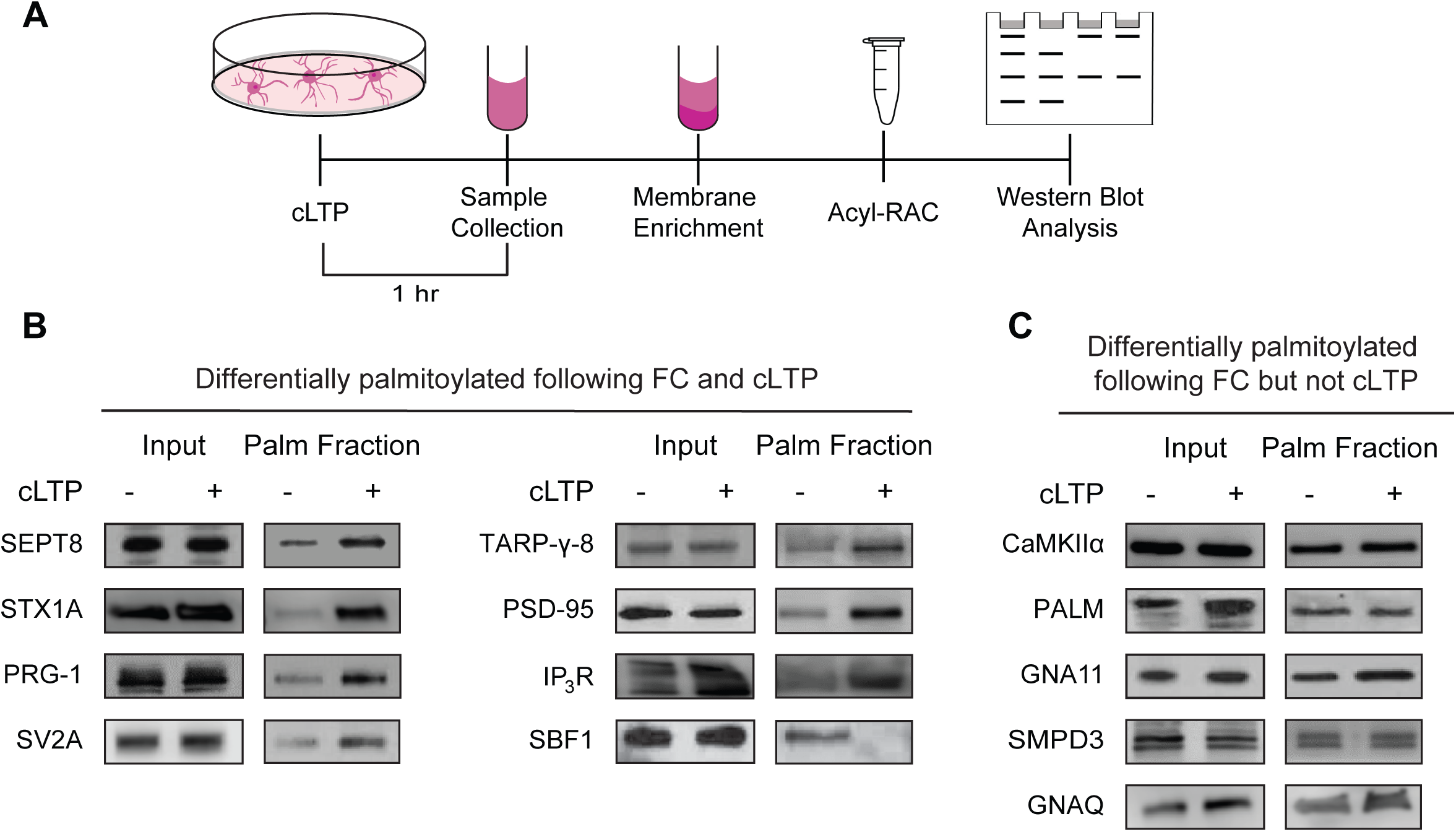
Validation of differentially palmitoylated proteins. **(A)** Schematic illustration of the *in vitro* assay to validate activity-induced changes in neuronal proteins. **(B and C)** Acyl RAC assays demonstrate that a subset of proteins that are differentially palmitoylated in response to FC (*p* <0.05 and/or fold enrichment ≥ 2), also undergo dynamic palmitoylation 1 hour post cLTP induction **(B)**, whereas a subset of proteins did not **(C)**. 7 of the 8 proteins that are differentially palmitoylated exhibit increased palmitoylation after both FC and cLTP, whereas SBF1 exhibits decreased palmitoylation after both FC and cLTP.

### Activity-dependent palmitoylation of postsynaptic protein PRG-1

We next wanted to investigate the downstream effects of activity-induced changes in palmitoylation on synaptic plasticity. We focused on PRG-1, a palmitoylation substrate identified in our proteomic screen that has already been shown to play a significant role in regulating synapse plasticity in the hippocampus (Liu et al., 2016). PRG-1 belongs to a family of transmembrane proteins with structural homology to the lipid phosphate phosphatases (LPPs) and thus are referred to as lipid phosphate phosphatase-related proteins (LPPRs), or plasticity-related genes (PRGs), which instead reflects their role in neuronal plasticity (Sigal, McDermott, & Morris, 2005). We demonstrated that PRG-1 palmitoylation significantly increases 10 minutes following cLTP treatment (1.4 ± 0.1-fold) and remains elevated for 1 hr (1.7 ± 0.2-fold) (Fig. 5, A and B). There are three known isoforms of human PRG-1 corresponding to 62, 77, and 83 kDa in molecular weight (https://www.uniprot.org/uniprot/Q7Z2D5). The same alternative splicing sites are also predicted for the rat PRG-1 protein, which would produce protein isoforms of similar sizes. Using the acyl-RAC assay, palmitoylation was predominantly detected for the 83 kDa isoform of PRG-1, suggesting that this is the main palmitoylated form of the protein (Fig. 5 A). Palmitoylation of transmembrane proteins typically occurs on cysteine residues adjacent to or within the transmembrane domain (Blaskovic, Blanc, & Van Der Goot, 2013). In line with this, the palmitoylation site prediction software CSS-Palm 4.0 (Zhou, Xue, Yao, & Xu, 2006) (http://csspalm.biocuckoo.org/) revealed two cysteine residues (C146 and C147 in human; C147 and C148 in rat) in a juxtamembrane region of PRG-1, facing the cytosolic side of the membrane.

**Figure 5:**
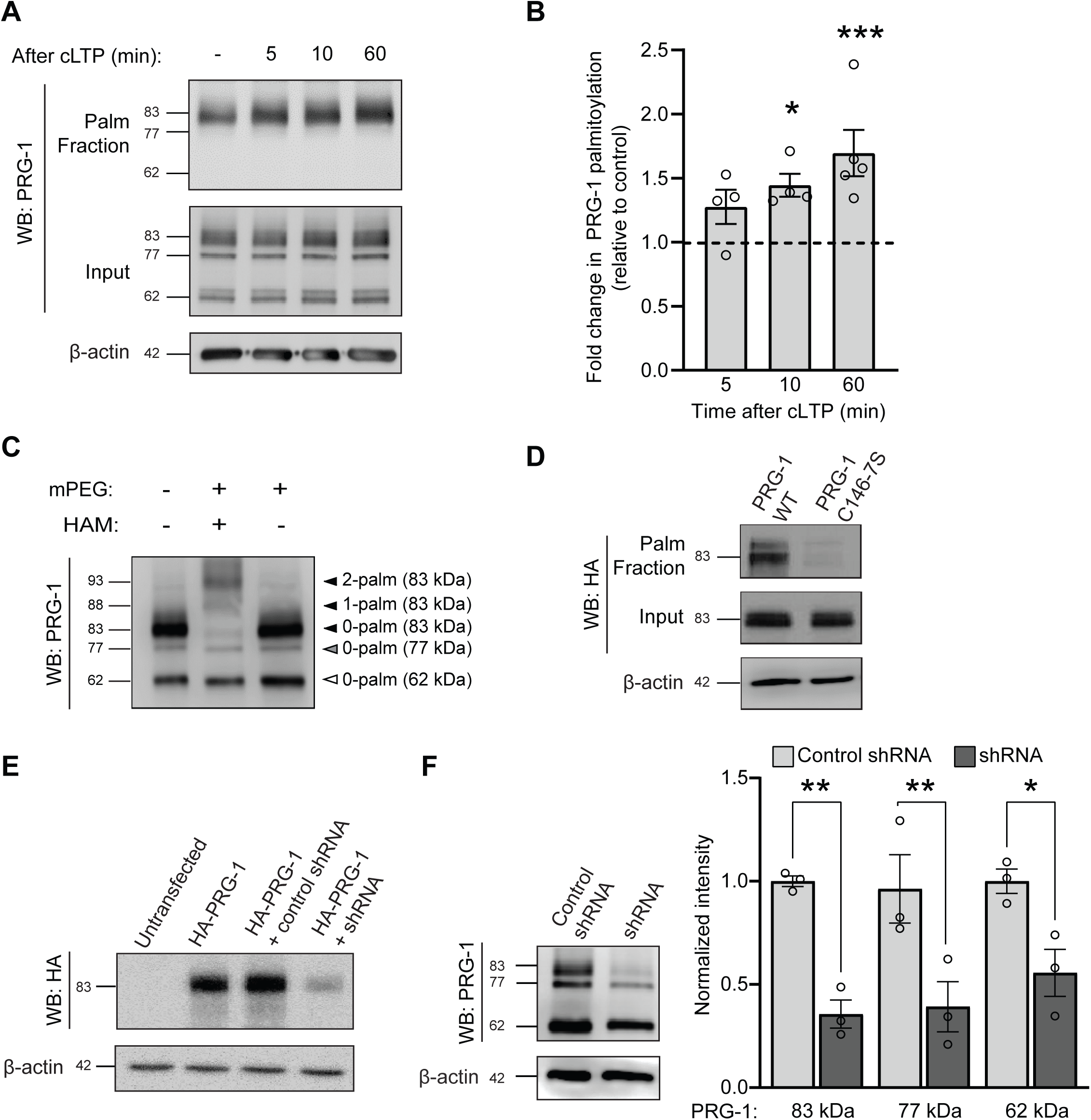
Palmitoylation of PRG-1 in response to synaptic activity in hippocampal neurons. **(A and B)** Acyl-RAC assay demonstrating a significant increase in PRG-1 palmitoylation 10 minutes and 1 hour following cLTP induction relative to mock control. Only the 83kDa isoform is palmitoylated. *n* = 4 blots from 4 cultures for 5 and 10 min cLTP; *n* = 5 blots from 5 cultures for 1 hour. *p < 0.05, ***p < 0.001, one-way ANOVA, Tukey’s test *post-hoc*. **(C)** PRG1 is palmitoylated on two cysteine residues in hippocampal neurons. APEGs assay depicting the number of palmitoylation-dependent mobility shifts of PRG-1 1 hour after mock or cLTP stimulation. *n* = 3 blots from 3 cultures. **(D)** Palmitoylation of PRG-1 occurs at cysteines 146 and 147. 0 DIV hippocampal neurons were nucleofected with HA-PRG-1 WT or HA-PRG-1 C146-7S mutant constructs, followed by an acyl-RAC assay /western blot at 7 DIV. Palmitoylation of the PRG-1 C146-7S mutant was greatly reduced compared to WT PRG-1. **(E and F)** shRNA-mediated knockdown of PRG1 in HEK293T cells **(E)** and primary hippocampal neurons **(F)**. **(E)** HEK cells were either untransfected or transfected with HA-PRG-1 alone or in combination with control shRNA or PRG-1 shRNA. Blots were probed with anti-HA antibody. *n* = 3 blots from 3 cultures. **(F)** Primary hippocampal neurons were nucleofected with control or PRG-1 shRNA at 0 DIV and blots probed with anti-PRG-1 antibody. There was a significant reduction in all 3 isoforms. *n* = 3 blots, 3 cultures. \**p* < 0.05; \*\**p* < 0.01 relative to control shRNA, two-way ANOVA, Bonferroni’s test *post-hoc.* Mean ± SEM for **(B)** and **(F)**.

To quantify the number of palmitoylated residues and to determine which residues are specifically palmitoylated following increased synaptic activity, cLTP was induced in 14 DIV cultured hippocampal neurons followed by an Acyl PEGyl Gel Shift (APEGS) assay 1 hr later. When proteins are separated by SDS-PAGE in this protocol, labeling of palmitoylated cysteines with mPEG-5k causes a mobility shift of 5 kDa (Yokoi et al., 2016). The APEGS assay revealed two distinct mobility shifts from the endogenous PRG-1 protein sizes after cLTP, and this mobility shift was again primarily observed in the 83 kDa isoform (Fig. 5 C). Moving forward, the palmitoylation of this specific isoform was investigated. Cysteine residues, C146 and 147, on the 83 kDa isoform of PRG-1 that were predicted to be palmitoylated were then mutated to serines using site-directed mutagenesis (PRG-1 C146-7S). The robust palmitoylation observed for wildtype HA-tagged PRG-1 (PRG-1 WT) was abolished in neurons expressing HA-tagged PRG-1 C146-7S mutant (Fig. 5 D), revealing that these residues are the sites of PRG-1 palmitoylation.

### PRG-1 palmitoylation is required for AMPAR insertion following cLTP treatment

To determine whether activity-induced palmitoylation of PRG-1 is important for the regulation of synapse plasticity, we tested whether the palmitoylation of PRG-1 impacted the incorporation of GluA1-containing AMPARs into the postsynaptic membrane. To knockdown endogenous PRG-1, we used a short hairpin RNA (shRNA) that we validated in HEK293T cells and primary hippocampal neurons (Fig. 5, E and F, respectively). In neurons, PRG-1 shRNA was shown to knockdown all 3 endogenous PRG-1 isoforms, the palmitoylated 83kDa isoform as well as the unpalmitoylated 77 and 62 kDa isoforms (Fig. 5 F).

To investigate if PRG-1 palmitoylation is required for activity-induced AMPAR insertion, we tested cLTP induced plasma membrane insertion of the AMPAR GluA1 subunit tagged with superecliptic pHluorin (SEP), a pH-sensitive variant of green fluorescent protein that has previously been used to evaluate the strengthening and weakening of synapses (Ashby et al., 2004; Brigidi et al., 2015; Makino & Malinow, 2009; McLeod et al., 2018; Miesenböck, De Angelis, & Rothman, 1998; Patterson, Szatmari, & Yasuda, 2010). Cells were co-transfected with an mCherry cell fill, SEP-GluA1 and either a control shRNA, PRG-1 shRNA, or PRG-1 shRNA plus either an shRNA resistant PRG-1 (PRG-1 WT^R^) or shRNA resistant and palmitoylation-deficient PRG-1 (PRG-1 C146-7S^R^), as indicated (Fig. 6). We observed a decrease in the basal density and intensity of SEP-GluA1 puncta with both PRG-1 shRNA or shRNA plus PRG-1 C146-7S^R^ (Fig. 6 A-C). As previously reported (Brigidi et al., 2015), we observed a significant increase in the intensity of SEP-GluA1 clusters 20 min after cLTP stimulation in control cells or those expressing PRG-1 shRNA plus PRG-1 WT^R^ (Fig. 6, A and D). In contrast, PRG-1 knockdown or loss of PRG-1 palmitoylation abrogated the activity-induced increase in SEP-GluA1 intensity (Fig. 6 D). Together, these results suggest that PRG-1 is involved in activity-induced recruitment of AMPARs, and that PRG-1 palmitoylation is essential for this function.

**Figure 6:**
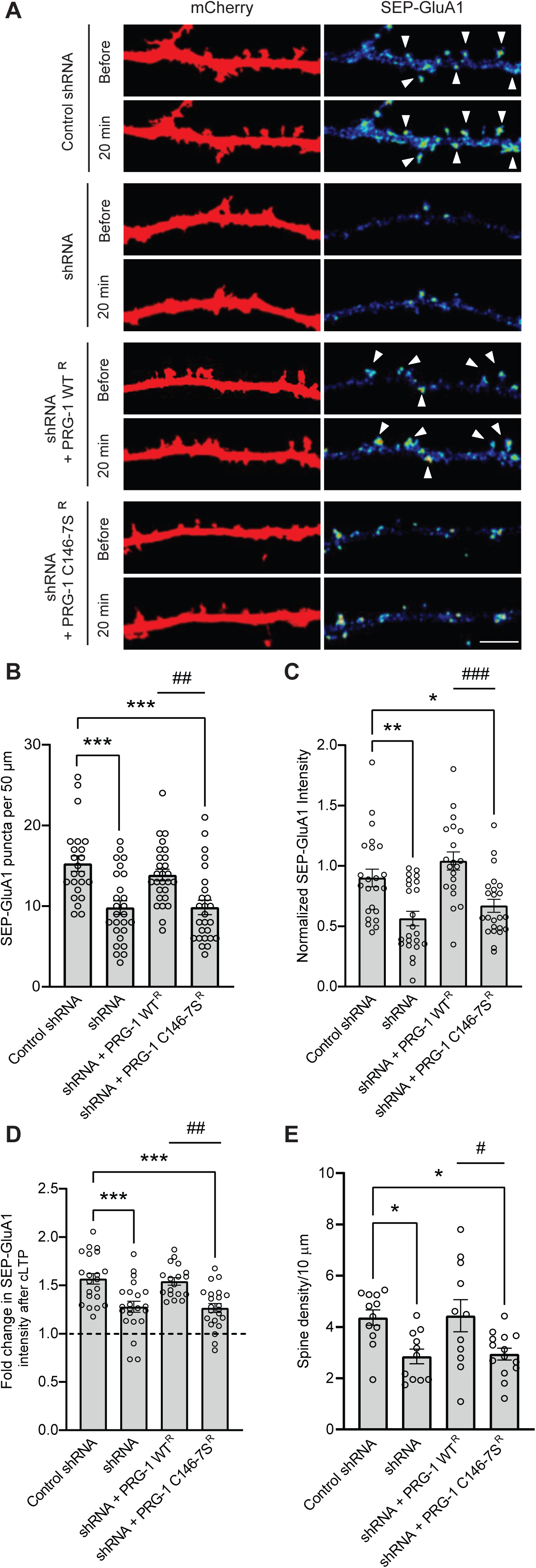
PRG-1 palmitoylation is important for activity-induced insertion of AMPAR into postsynaptic membranes. **(A)** Confocal images of 14 DIV hippocampal neurons transfected at 11 DIV with SEP-GluA1, mCherry, and indicated constructs before and 20 min after glycine stimulation. SEP-fluorescent puncta are pseudocolored as heat maps. Scale bar, 5 μm. The density of both spines **(A and E)** and SEP-GluA1 puncta **(A and B)** are significantly reduced in PRG-1 knockdown cells and those expressing PRG-1 palmitoylation-defective mutant (PRG-1 C146-7S^R^) compared to controls. The intensity of SEP-GluA1 puncta is also reduced in PRG-1 knockdown cells and those expressing PRG-1 C146-7S^R^ mutant **(A and C)**, and cLTP-mediated increase in surface GluA1 levels is abolished in PRG-1 knockdown cells and those expressing the palmitoylation-defective PRG-1 C146-7S^R^ mutant. **(A and D)**. *n* = 23 control, *n* = 26 shRNA, *n* = 27 shRNA + PRG-1^R^, *n* = 26 shRNA + C146-7S^R^ neurons from 3 cultures for **(B)**. *n* = 23 control, *n* = 23 shRNA, *n* = 20 shRNA + PRG-1^R^, *n* = 23 shRNA + C146-7S^R^ neurons from 3 cultures for **(C)**. *n* = 21 control, *n* = 22 shRNA, *n* = 18 shRNA + PRG-1^R^, *n* = 22 shRNA + C146-7S^R^ neurons from 3 cultures for **(D)**. n = 12 control, 11 shRNA, 11 shRNA + PRG-1^R^, 14 shRNA + C146-7S^R^ neurons from 11 cultures for **(E)**. \**p* < 0.05, \*\**p* < 0.01, \*\*\**p* < 0.1 relative to control; #*p* < 0.05, ##*p* < 0.01, ###*p* < 0.001 relative to shRNA + PRG-1^R^, one-way ANOVA, Tukey’s test *post-hoc*. Graph displays mean ± SEM for **(B-E)**.

We also observed a significant decrease in the density of postsynaptic spines following PRG-1 knockdown (Fig. 6, A and E) in accordance with published work showing that PRG-1 regulates hippocampal spine formation and density *in vivo* and *in vitro* (Liu et al., 2016). While PRG-1 WT^R^ rescued the spine density decrease, palmitoylation-defective PRG-1 C146-7S^R^ did not, indicating that palmitoylation of PRG-1 is important in spine formation and/or maintenance. Together, these results demonstrate that the palmitoylation of PRG-1 can impact both basal synapse density and activity-dependent changes in AMPAR trafficking that are critical for the consolidation of LTP.

Palmitoylation of PRG-1 may impact its function by altering its subcellular localization. Using surface biotinylation assays (Fig. 7 A-C) and immunocytochemical colocalization analysis (Fig. 7, D and E), we demonstrated that synaptic activity does not impact PRG-1 surface expression nor its localization at synapses. This suggests that activity-induced palmitoylation of PRG-1 impacts PRG-1 function and that this does not depend on PRG-1 membrane trafficking.

**Figure 7:**
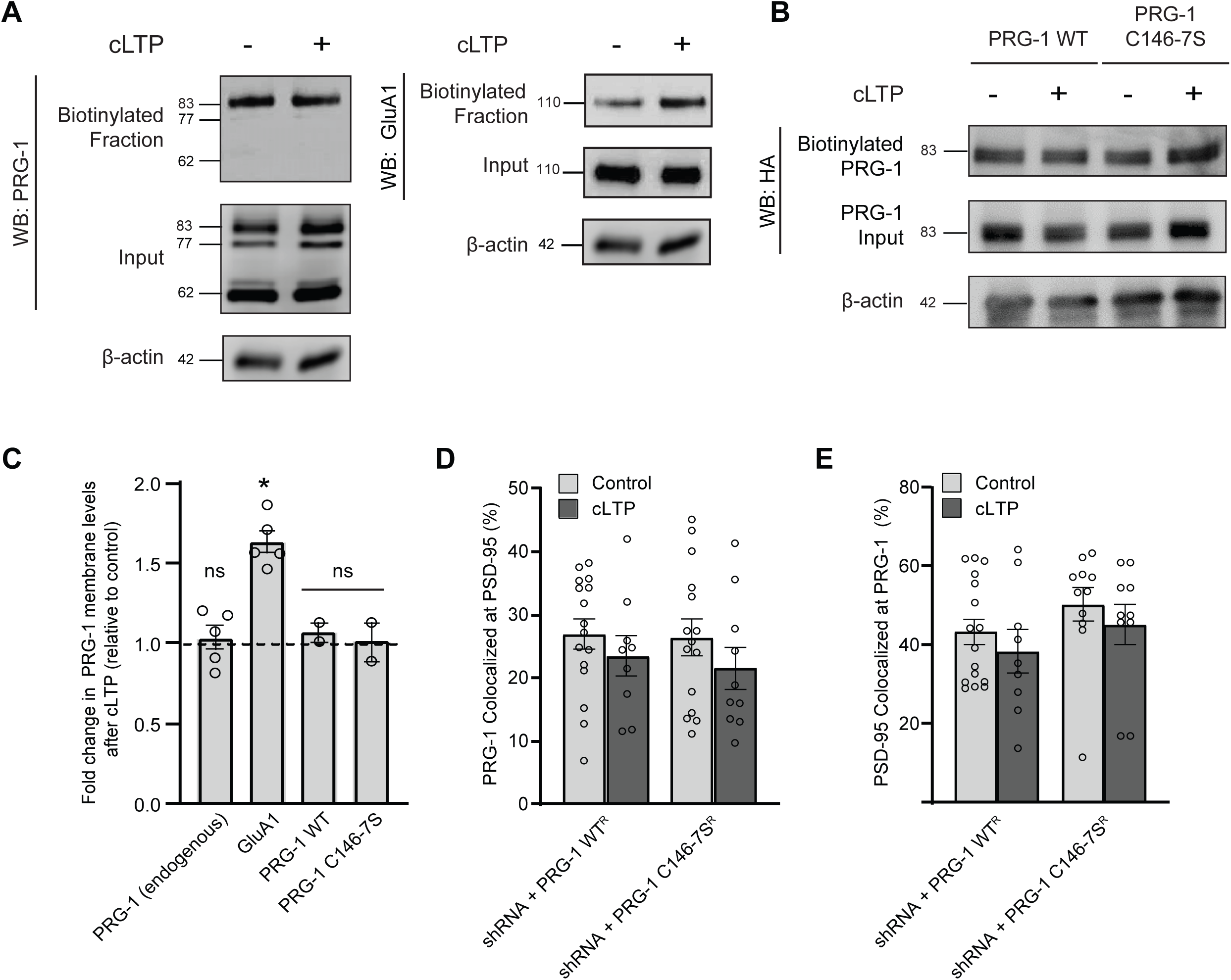
PRG-1 palmitoylation does not regulate its membrane expression or synaptic localization. **(A and B)** Biotinylation assays in 14 DIV hippocampal cultures 1 hour after treatment with glycine (+cLTP) or control buffer (-cLTP). Surface membrane proteins were isolated by immunoprecipitating with neutravidin-coated beads prior to immunoblotting. **(A and C)** There is no difference in endogenous surface PRG-1 levels 1 hr following cLTP. In contrast, endogenous GluA1, used as a control for the efficacy of the cLTP, was robustly recruited to the cell surface. **(B and C)** Disrupting the palmitoylation of PRG-1 (PRG-1 C146-7S) does not impact its basal surface expression compared to PRG-1 WT and neither PRG-1 WT nor PRG-1 C146-7S exhibit activity-induced changes in surface expression. *n* = 5 blots from 5 cultures, \**p* < 0.05 relative to control surface levels for each protein, one-way ANOVA. **(D and E)** 15 DIV hippocampal neurons transfected with the indicated constructs and immunostained for PRG-1 and the postsynaptic marker PSD-95 demonstrate no change in the postsynaptic localization of PRG-1 1 hr after cLTP induction as indicated by colocalization. *n* = 9-16 cells, from 3 cultures. Graph displays mean ± SEM for **(C-E)**.

## DISCUSSION

This study demonstrates that contextual fear conditioning regulates the palmitoylation of a large number of proteins in the hippocampus, the majority of which are synaptic proteins shown to play a role in regulating synapse formation and plasticity. GO and STRING analyses revealed two main groups of substrates that are differentially palmitoylated following fear conditioning – a group of non-synaptic, metabolic proteins which predominantly exhibit decreased palmitoylation in fear-conditioned mice, and a second group of synaptic proteins which predominantly exhibit increased palmitoylation levels in the fear-conditioned group relative to control. We demonstrate that dynamically palmitoylated proteins are highly co-expressed in neurons together with a specific subset of palmitoylating enzymes and co-factors, enabling the generation of multiple hypotheses regarding enzyme/substrate pairs. The terms most overrepresented in our gene list included ones associated with synapse plasticity such as “synaptic vesicle cycle”, “calcium signalling pathway” and “glutamatergic synapse”, underscoring the role of dynamic palmitoylation in synapse plasticity. Exploring this further, we demonstrate that palmitoylation of the synaptic protein, PRG-1, is increased in response to both hippocampal-dependent contextual fear conditioning and cLTP. We demonstrate that this activity-mediated palmitoylation is important for the insertion of AMPARs into the synaptic membrane, which is thought to be essential for increases in synapse strength (Petrini et al., 2009). Together, these findings are consistent with a critical role for regulated palmitoylation in modulating changes in synaptic transmission and plasticity that underlie hippocampal learning and memory processes.

We found that changes in protein palmitoylation after FC are not secondary to changes in total protein turnover. This is in accordance with a prior study which demonstrated changes in the mouse hippocampal synaptic membrane proteome 4 hours, but not 1 hour, following contextual fear learning (Rao-Ruiz et al., 2015). Similarly, a comparison of our list of differentially palmitoylated proteins with those that were differentially palmitoylated in response to chronic restraint stress followed by tail suspension stress (Zareba-Koziol et al., 2019) revealed only 5 overlapping proteins, providing additional confidence that our list represents proteins that are dynamically palmitoylated in response to the contextual learning associated with fear conditioning rather than associated stress. Previous work has examined activity-dependent regulation of candidate hippocampal proteins of interest using various protocols, while a more unbiased assessment of all proteins that can be dynamically palmitoylated in response to a more physiological learning paradigm has not been explored. Kang and colleagues (Kang et al., 2008) examined changes in the palmitoylation of candidate synaptic proteins using a kainic acid seizure model in rats. Five of the nine candidate proteins found to be differentially palmitoylated in response to kainic acid treatment in that study were also identified within our list of 121 substrates.

When examining gene expression using the DropViz hippocampal single-cell RNAseq database, we found that 10 of the 30 palmitoylating/ de-palmitoylating enzymes and accessory proteins were predominantly enriched in neurons and also highly co-expressed with neuron-enriched differentially palmitoylated proteins from our screen. ZDHHC enzymes in this subset are thought to localize to a variety of subcellular domains, including the Golgi (ZDHHC3, ZDHHC13, ZDHHC17, ZDHHC21, ZDHHC23) and postsynapse (ZDHHC2, ZDHHC5, ZDHHC8) (Malgapo & Linder, 2021; Solis, Valnohova, Alvarez, & Katanaev, 2020). We found that a substantial proportion of the differentially palmitoylated substrates identified in our screen localize at the pre- and/or postsynapse, making ZDHHC2, ZDHHC5 and ZDHHC8 well positioned to mediate postsynaptic increases in substrate palmitoylation in response to plasticity-inducing stimuli. Accordingly, ZDHHC2 and ZDHHC5 have been found to dynamically palmitoylate their substrates in response to synaptic activity (Fukata et al., 2013; Woolfrey et al., 2015). However, it is less clear which ZDHHC enzymes might mediate differential palmitoylation of presynaptic substrates. Indeed, while ZDHHC5 and ZDHHC8 were found to localize to axons in peripheral dorsal root ganglia neurons (Collura et al., 2020), reports of ZDHHCs localizing to the pre-synapse in the central neurons of the brain are currently limited. Although several presynaptic palmitoylation substrates have been identified previously, including SNAP25 (Greaves, Gorleku, Salaun, & Chamberlain, 2010) and GAD65 (Verardi, Kim, Ghirlando, & Banerjee, 2017), we report here a novel extensive list of presynaptic proteins that are dynamically palmitoylated in response to a learning event. Differential palmitoylation is therefore likely to have considerable influence over numerous aspects of presynaptic function. An important goal of future research will be to identify any ZDHHCs localized to the presynapse that may these effects in the hippocampus.

We also observed differential palmitoylation in non-neuronal enriched genes such as *Cnp* and *Plp* that are highly expressed in oligodendrocytes, indicating a possible role for differential palmitoylation in activity-dependent myelin remodeling in the hippocampus (Gibson et al., 2014; Steadman et al., 2020). In addition, we found differential palmitoylation of astrocyte enriched substrates including SLC1A2 and GPR37L1, which may alter the function of the tripartite synapse (Jolly et al., 2018; Kugler & Schleyer, 2004). While the role of palmitoylation is relatively understudied in non-neuronal cell types of the brain, our data suggest that activity-dependent palmitoylation may also be important for regulation of glial cell function.

Our results demonstrate that only one of the three rat PRG-1 isoforms (which correspond to isoforms found in humans) is palmitoylated *in vitro*. This likely reflects strong substrate specificity by ZDHHC enzymes as the dual palmitoylation site is found in the protein sequence of all three PRG-1 splice forms. Differential palmitoylation of splice variants has been previously reported for other proteins localized at the synapse, including the brain-specific isoform of Cdc42 (Kang et al., 2008) as well as the SNAP25a and SNAP25b isoforms (Greaves et al., 2010), which may reflect unique regulation of alternatively spliced variants in neurons. Interestingly, our results also indicate that the palmitoylated 83 kDa isoform of PRG-1 is most abundantly localized to the surface plasma membrane, compared to the other variants. Together, these findings may suggest that the canonical PRG-1 isoform has a distinct function that requires both precise modulation by changes in palmitoylation and enriched membrane abundance.

Prior work has demonstrated a reduction in postsynaptic spine density and impairments in LTP in PRG-1 knockout mice (Liu et al., 2016). PRG-1 is thought to regulate spine formation and synapse plasticity by interacting with protein phosphatase 2A (PP2A) which in turn recruits and activates β1-integrin (Liu et al., 2016). PRG-1 regulation of synapse plasticity has also been shown to rely on a non-cell autonomous mechanism involving PRG-1-mediated removal of lysophosphatidic acid (LPA) from the synaptic cleft and the subsequent control of LPA-mediated glutamate release via presynaptic LPA receptors (Trimbuch et al., 2009). Our observation that palmitoylation is important for both spine formation and the recruitment of AMPARs postsynaptically suggests that PRG-1 palmitoylation controls either the binding to PP2A, its ability to bind and clear LPA, or both. Additional work is required to tease out exact mechanisms of action downstream of PRG-1 palmitoylation.

Regulated palmitoylation may play a role in modulating the changes in synaptic transmission and plasticity that underlie hippocampal memory processes. Our analysis of the subcellular localization of differentially palmitoylated proteins revealed a broad distribution throughout major pre- and postsynaptic compartments, including synaptic vesicle components, synapse adhesions and the postsynaptic density, as well as a few astrocyte-enriched proteins, completing the tripartite synapse. As the hippocampus is considered to encode context representations (Maren, Phan, & Liberzon, 2013), it is possible that the enhancement or suppression of palmitate turnover on specific hippocampal proteins contributes to the context encoding aspect of associative fear learning. These novel networks of dynamically palmitoylated proteins may play a central role in learning and memory processes under physiological conditions, which may be disrupted in brain pathology.

## METHODS

### Antibodies

Primary antibodies used were as follows: **Calcium Voltage-Gated Channel Auxiliary Subunit Gamma 8** (1:500; BD Transduction Laboratories 610921), **Calnexin** (1:500; Life Technologies MA5-13134), **CamKIIa** (1:1,000; Life Technologies MA1-16746), **CamKv** (1:500; BD Transduction Laboratories 610163), **CRMP4** (1:500; Millipore AB5905), **Guanine nucleotide-binding protein subunit alpha-11** (1:1,000; Millipore 05-855R), **Guanine nucleotide-binding protein G(q) subunit alpha** (1:1,000; AbCam ab75825)**, HA** (1:1,000 (WB); Cell Signalling Technology #3724; 1:1,000 (ICC); Sigma H9658), **Paralemmin** (1:1,000, Acris-OriGene TA335984), **Plasticity related protein-1** (1:1,000 (WB); Abcam ab104104), **PSD-95** (1:1,000 (WB); Calbiochem CP35; 1:500 (ICC); Abcam ab2723), **Sbf1** (1:1,000; Covance MMS-101P), **Septin 8** (1:500; Abnova H000055737-M02), **Slc32a1** (1:1,000; Roche 11814460001), **Smpd3** (1:500; Cell Signaling Technology C29F4), **Synaptic vesicle protein 2A** (1:500; Millipore GR09L), **Syntaxin-1a** (1:500; Synaptic Systems 132 002), **VGLUT1** (1:1,000; Cell Signaling Technologies 2102S). IgG-horseradish peroxidase-conjugated antibodies were obtained from Bio-Rad (Hercules, CA): **goat anti-mouse** (1:5,000; 170-6516) and **goat anti-rabbit** (1:5,000; 170-6515). Fluorescent secondary antibodies used were: **Alexa-Fluor 405 goat anti-guinea pig** (1:1,000; Abcam ab175678), **Alexa-Fluor 568 goat anti-mouse IgG2a** (1:1,000; Life Technologies A-21134), **Alexa-Fluor 568 goat anti-mouse IgG1** (1:1,000; Life Technologies A-21124) and **Alexa-Fluor 633 goat anti-rabbit** (1:1,000; Life Technologies A-21070).

### Plasmids and Primers

The rat HA-tagged PRG-1 ORF construct was generated by Gibson Assembly Cloning from NEB (Ipswich, MA, #E5510S) of the Plppr4 ORF clone (NM_001001508.2) from GenScript (Piscataway, NJ, #SC1200) to introduce an N-terminal 3xHA tag 5’-ACCCATACGATGTTCCAGATTACGCTTACCCATACGATGTTCCAGATTACGCTTACCCATACGATGTTCCAGATTACGCT-3’. The shRNA target sequence that gave maximum knockdown efficiency of rat PRG-1 was 5′-GCAAGAACGAGAGTCGCAAGA-3′ and was obtained from GeneCopoeia (Rockville, MD, #RSH090356). Human PRG-1 obtained from DNASU Plasmid Repository (The Biodesign Institute/Arizona State University, AZ, #HsCD00080375), was not targeted by the shRNA and used for rescue experiments (PRG-1 WT^R^). To generate PRG-1 that cannot be palmitoylated (PRG-1 C146-7S^R^), site-directed mutagenesis (Agilent Technologies, Santa Clara, CA: QuikChange II XL Site-Directed Mutagenesis Kit, 200521) of the human PRG-1 ORF sequence AGAAGGAATTCTCTAC<u>AGTAGC</u>CTCTCCAAAAGAAGAAATGGGGTC was done using the following primers: Forward 5′-GGCGAAGGAGAGGCCAAAGG-3′ and Reverse 5′-AACTGTGAGGTCAGATCCTGAGC-3′ (underlined residues indicate amino acid codons where nucleotide bases were mutated). The SEP-GluA1 construct was generated by Dr Roberto Malinow (University of California, San Diego) and obtained from Addgene (Watertown, MA: Plasmid #24000).

### Primary Hippocampal Cultures

Hippocampi were harvested from embryonic day 18 (E18) Sprague–Dawley rats of either sex as previously described and plated onto plastic 6-well plates at a density of 130 cells per mm^2^ or on 18 mm coverslips (Marienfeld, Lauda-Königshofen, Germany) (Shah et al., 2019).

### Transfection (primary hippocampal cultures/HEK293T cells)

Primary hippocampal cultures were transfected at 9–10 DIV by using Lipofectamine 2000 (Invitrogen, Carlsbad, CA) according to the manufacturer’s recommendations. Cultured neurons plated on coverslips were then live imaged (14 DIV) for subsequent experiments. HEK293T cells were transfected with Lipofectamine 2000 (Invitrogen) at 70% confluency at a ratio of 3:1 (shRNA or scrambled shRNA to target plasmid) and incubated for 70–72 h before harvesting for biochemical analysis. For nucleofection, hippocampi were dissociated as above and ∼6 million neurons were isolated and plated onto a 10-cm dish for future biochemical experiments. Neuronal nucleofections were performed immediately before plating at 0 DIV by using an Amaxa Nucleofection Kit (Lonza, Basel, Switzerland, VPG-1003) according to the manufacturer’s optimized protocol (Number 101, program G-13), and were used for experiments at either 6-7 DIV (shRNA validation experiments) or 14 DIV (acyl-RAC and APEGS assays).

### Animals

9-week-old, male C57BL/6 mice were used for fear conditioning experiments. Mice were purchased from the Jackson Laboratory and group-housed in the University of British Columbia animal facility. All procedures involving animals were approved by the Canadian Council of Animal Care and the University of British Columbia Committee on Animal Care.

### Context- dependent Fear Conditioning

Mice were habituated to handling and to the environmental stimuli of the experimental room for 3 min/day for 6 consecutive days (Radulovic, Kammermeier, & Spiess, 1998). Mice were placed in the behavior room for 30 min before the experiment. Context-dependent fear conditioning experiments were performed in a training cage equipped with stainless-steel shocking grids connected to a precision feedback current-regulated shocker (Med Associates, Inc., St. Albans, VT, USA). The chamber was cleaned with 70% ethanol prior to each session. The mice were placed into the fear conditioning training box in a randomized order and allowed to freely explore the apparatus for 3 min. After 3 minutes the conditioned group (*n* = 8 mice) received a foot shock stimulus (0.3 mA, 50 Hz) lasting 5 seconds whereas the control group (*n* = 8 mice) received no foot shock (Brigidi et al., 2014). Mice were returned to their home cages five minutes after conditioning. 1 hour later, context-dependent freezing was measured. Mice were placed in the same illuminated compartment and observed for the presence/absence of a freezing response over 5 minutes(Button et al., 2019). During this time, the mice were not exposed to any shock and the percent freezing time was measured using the ANY-Maze system (Stoelting Co., Wood Dale, IL). Freezing behavior was video-tracked and analyzed on a second-by-second basis and defined as the absence of movement except for respiration. Experiments were performed during the dark (wake) cycle.

### Liquid Chromatography-Tandem Mass Spectrometry (LC-MS/MS)

Protein precipitates, boiled in SDS sample buffer, were run on 10% (wt/vol) SDS/PAGE gel. Proteins were visualized by colloidal Coomassie and digested out of the gel as described previously (Chan, Howes, & Foster, 2006). Samples were purified by solid phase extraction on C-18 STop And Go Extraction (STAGE) Tips. Following purification, samples were analyzed by a quadrupole–time of flight mass spectrometer (Impact II; Bruker Daltonics, Billerica, MA) coupled to an Easy nano LC 1000 HPLC (Thermo Fisher Scientific, Waltham, MA). The analytical column used was 40–50 cm long, with a 75-μm inner diameter fused silica with an integrated spray tip pulled with P-2000 laser puller (Sutter Instruments, Novato, CA) and packed with 1.9 μm diameter Reprosil-Pur C-18-AQ beads (Dr. Maisch, Ammerbuch, DE). Buffer A consisted of 0.1% aqueous formic acid, and buffer B consisted of 0.1% formic acid and 80% (vol/vol) acetonitrile in water. After a standard 90-min peptide separation, the column was washed with 100% buffer B before re-equilibration with buffer A. The Impact II was set to acquire in a data-dependent auto-MS/MS mode with inactive focus fragmenting the 20 most abundant ions (one at the time at 18-Hz rate) after each full-range scan from *m/z* 200 to *m/z* 2,000 at 5 Hz rate. The isolation window used varied between 2–3 depending on the parent ion mass to charge ratio, and the collision energy ranged from 23 to 65 eV depending on ion mass and charge. Parent ions were excluded from MS/MS for the first 0.4 min and revaluated only if their intensity increased more than five times. Singly charged ions were excluded from fragmentation.

### Protein Search and Analysis

Data was analyzed with MaxQuant 1.5.3.30 (Cox & Mann, 2008; Tyanova, Temu, & Cox, 2016) and quantitation performed using peptide intensities with match between runs enabled(Collins, Woodley, & Choudhary, 2017; Meier, Geyer, Virreira Winter, Cox, & Mann, 2018). The searches with the MaxQuant algorithm were matched to a *mus musculus* database downloaded from uniprot.org (reviewed and unreviewed; RRID:SCR_002380) and a contaminant and decoy database. Default search parameters were used except for the following: a 0.15 Da first search and 0.008 Da main search tolerance, fixed carbamidomethyl (C) modification, MSMS match tolerance of 20 ppm, and MSMS not required for LFQ. Accurately identified peptides were only confined to those with IonScores higher than 99% confidence. Contaminant and reverse hits were removed. In LFQ, the intensities of the thus recorded peaks were taken as proxies for peptide abundance. To identify the peaks observed in the MS1 spectrum, these peaks were isolated and fragmented.

The resulting fragmentation spectra (so-called MS2 spectra) was then used for peptide identification. In LFQ, each sample was separately analyzed on the mass spectrometer, and differential expression was obtained by comparing relative intensities between runs for the same identified peptide (Goeminne, Gevaert, & Clement, 2018). For quantification, LFQ protein amount values were log_2_-transformed, to reduce the effect of outliers, prior to hypothesis testing (t-test) (*p* < 0.05). Log ratios were calculated as the difference in average log_2_ LFQ intensity values between control and conditioned groups. Two-tailed, Student’s t test calculations were used in statistical analysis. For each protein with sufficient data, three significance values were calculated between three pairs of conditions:

FC (+)HAM vs Control (+)HAM, FC (+)HAM vs FC (-)HAM, and Control (+)HAM vs Control (-)HAM. Proteins were deemed to have sufficient data for a hypothesis test if they had at least two valid values in both conditions being compared. Reproducibility of protein quantitation was also assessed using pairwise replicate-to-replicate correlations (Pearson correlation). Proteins with reproducible quantitation had higher Pearson correlation values across pairs of replicates.

### Bioinformatics

SwissPalm (2019-09-08) (Blanc et al., 2015) was used to determine which of the 345 palmitoylated proteins quantified were previously indexed as predicted or validated substrates for palmitoylation. The Mouse Genome Informatics (RRID:SCR_006460) database(Bult et al., 2019) was used to assign GO classifications for the list of 121 differentially palmitoylated proteins. KEGG pathway analysis was performed using the web-based program g:Profiler (Raudvere et al., 2019) and the mouse genome reference data set, with a *p*-value threshold of 0.05 (Raudvere et al., 2019). DiGeNET (Piñero et al., 2019) disease enrichment analysis was conducted by a multigene query in EnrichR (Chen et al., 2013; Kuleshov et al., 2016). Functional interaction networks of the 121 differentially palmitoylated proteins were investigated using the Search Tool for the Retrieval of Interacting Genes (STRING) 11.0 (Szklarczyk et al., 2019). A threshold confidence level of 0.4 was used to identify protein interactions and seven types of protein interactions were used for network generation, including neighborhood, gene fusion, co-occurrence, co-expression, experimental, database knowledge, and text mining. Co-expression analysis (Spearman correlation) of the 121 differentially palmitoylated proteins with 30 known palmitoylating/depalmitoylating enzymes and accessory proteins was performed across the 103 transcriptionally unique cell populations in the mouse hippocampus (HC) database in DropViz. Correlation statistical analysis was performed using Hmisc R (https://cran.r-project.org/web/packages/Hmisc/index.html). Heatmaps of co-expression r-values and expression level z-scores were generated using the heatmaply package in R (Galili, O’Callaghan, Sidi, & Sievert, 2018). Enzymes and substrates were clustered by hierarchical clustering with a Euclidean distance metric and complete linkage (Fig. 3 A and B).

### Chemical LTP

14 DIV neurons were treated with a cLTP protocol described previously(Lu et al., 2001). Briefly, maintenance media was replaced with pre-warmed extracellular solution (ECS), supplemented with 0.5 μM tetrodotoxin, 1 μM strychnine, containing: 140 mM NaCl, 5 mM KCl, 1.3 mM CaCl_2_, 0 mM MgCl_2_, 25 mM HEPES, 33 mM D-glucose and a pH of 7.35, 290 mOsm l^−1^, for 10–15 min. For cLTP, this media was supplemented with 200 μM glycine (Sigma Aldrich, St. Louis, MO) for 3 min. After 3 mins, the cells were washed with ECS containing 2mM of MgCl_2_. The solution was then replaced with fresh Mg^2+^-free ECS for the indicated times before experimentation. Cells were maintained at 37 °C for the duration of activity stimulation.

### Preparation of Total Membrane Fraction

To prepare the total membrane fractions, ice cold homogenization buffer (20 mM HEPES, 1mM EDTA, 255mM sucrose, pH 7.4) with Complete Protease Inhibitors (Roche Diagnostics, Basel, CH) was added to samples and hippocampi were mechanically dissociated by passage through a 26-gauge syringe 10–15 times (Foster, De Hoog, & Mann, 2003). To clear nuclei, mitochondria, and large cellular debris, homogenates were centrifuged at 10,000 *g* for 10 min. The post nuclear supernatant was transferred to an ultracentrifuge tube and centrifuged for 2 h at 245,000 *g* at 4 °C in a Beckman L90 Ultracentrifuge (Beckman Coulter, Mississauga, Canada) to pellet remaining membrane.

### Acyl-RAC Assay

Membrane fractions were resuspended in blocking buffer containing methyl methanethiosulfonate (MMTS) to block the free thiols before incubation at 40°C for 4hr in a thermomixer. After incubation, 3 times the sample volume of cold acetone was added, and proteins were precipitated at −20°C for 20 min. Following centrifugation of the solution at 5,000 *g* for 10 min, the pellet was washed 5 times with 70% acetone, and air dried. The pellet was then resuspended in 300 μl of binding buffer (100 mM HEPES, 1.0 mM EDTA, 1% SDS, pH 7.5). Total protein was quantified with a bicinchoninic acid (BCA) assay (Pierce, Thermo Fisher Scientific, Waltham, MA) using bovine serum albumin (BSA) as the standard. Equal amounts of protein (0.5–2.0 mg for immunoblot experiments and 10–20 mg for mass spectrometry experiments) were diluted to a concentration of 2 mg/ml in binding buffer (100 mM HEPES, 1.0 mM EDTA, 1% SDS, pH 7.5) and added to ∼40 μl of prewashed thiopropyl Sepharose (Badrilla CAPTUREome™). To this mixture was added 19 μl of either 2 M HAM (NH_2_OH, freshly prepared in H_2_O from HCl salt and brought to pH 7.5 with concentrated NaOH) or 2 M NaCl. Binding reactions were carried out on a rotator at room temperature (RT) for 2.5 hr. Approximately 20 μl of each supernatant was saved as the “total input.” Resins were washed five times with binding buffer. For immunoblot analysis, elution was performed using 40 μl of SDS Laemmli sample buffer, heated to 70°C for 10 min, and separated via SDS-PAGE on a Mini-Gel apparatus (Bio-Rad, Hercules, CA).

### Western Blot Analysis

Western blotting was performed as previously described (Sun & Bamji, 2011a). Brain tissue and primary hippocampal neurons cells were homogenized in an ice-cold lysis buffer supplemented with phenylmethanesulfonyl fluoride solution and a protease inhibitor cocktail with EDTA (Roche). Whole-cell lysates were further homogenized by passage through a 26-gauge syringe 5–6 times. Lysates were cleared by centrifugation at 16,100*g* for 30 min at 4 °C and the solubilized fraction of protein was used for all biochemical experiments. Proteins were separated by SDS– PAGE (8-10% resolving gels, 4% stacking gels), analyzed by immunoblotting with the indicated antibodies and visualized using enhanced chemiluminescence (Pierce) on a Bio-Rad Versadoc 4000 (Bio-Rad).

### APEGS Assay

For the acyl-PEG assay, the commercially available SiteCounter™ palmitoylated protein kit (Badrilla, Leeds, UK) was used according to the manufacturer’s guidelines with the following optimizations: protein concentration was measured prior to the separation of experimental sample (Thioester Cleavage Reagent, +HAM) and negative control sample (Acyl Preservation Reagent, - HAM) using the BCA Assay (Pierce). Cultured hippocampal neurons were lysed and homogenized in blocking buffer (100 mM HEPES, 1.0 mM EDTA, 2.5% SDS, 0.1% MMTS, pH 7.5) to block the free thiols followed by incubation at 40°C for 4hr in a thermomixer. After incubation, three times the sample volume of cold acetone was added and then incubated at −20°C for 20 min to allow proteins to precipitate. Following the centrifugation of the solution at 16,000 *g* for 5 min, the protein pellet was washed 5 times with 70% acetone, and air dried. The pellet was then resuspended in 250 μl of binding buffer (100 mM HEPES, 1.0 mM EDTA, 1% SDS, pH 7.5). Total protein was quantified with the BCA assay (Pierce) using BSA as the standard. Resuspended proteins (0.4-0.75 mg) were incubated in binding buffer containing NH_2_OH (+HAM) for 1 hr at RT to cleave palmitoylated thioester bonds. As a negative control, an equal amount of protein was incubated with NaCl (-HAM). Following incubation, resuspended proteins were PEGylated for 1 hr at RT to label newly exposed cysteine thiols with the 5 kDa mass tag reagent (mPEG-5k, Badrilla SiteCounter™). Approximately 20 μl of each supernatant was saved as the “total input.” For immunoblot analysis, SDS Laemmli sample buffer was added to the protein samples, and then heated to 37°C for 30 min and separated via SDS-PAGE on a Mini-Gel apparatus (Bio-Rad).

### Immunocytochemistry

Immunocytochemistry experiments were performed as previously reported (Sun & Bamji, 2011b). Briefly, cells were fixed for 10 min in a pre-warmed (37°C) 4% paraformaldehyde–sucrose solution. Cells were then permeabilized in 0.1% Triton X-100 in PBS for 10 min at room temperature and blocked for 1 h at room temperature with 10% goat serum in PBS. Cells were then incubated in primary antibodies diluted in 1% goat serum in PBS overnight at 4°C. Cells were washed 3 times in PBS, incubated in secondary antibodies in 1% goat serum in PBS for 1 hr at RT. Cells were then washed 3 times in PBS and mounted on microscope slides using Prolong Gold (Molecular Probes, Thermo Fisher Scientific, Waltham, MA).

### Confocal Imaging

All fixed and live neurons were imaged using a Zeiss LSM 880 AxioObserver Airyscan microscope (Carl Zeiss Canada, North York, ON) with a Plan-Apochromat 63x/1.4 Oil DIC M27 objective. Identical acquisition parameters were used for all cells across all separate cultures within an experiment.

Images of fixed eGFP-expressing neurons were acquired using AiryScan Fast mode with a 0.55 μs dwell time (488, 543, and 633 nm lasers). The proximal dendritic arbor was imaged over a field of 132.9 μm x 132.9 μm at 0.4-μm intervals in the z-plane.

SEP–GluA1-expressing neurons were imaged in an artificial CSF (aCSF) imaging medium (137 mM NaCl, 2.7 mM KCl, 1 mM MgCl_2,_ 1.8 mM CaCl_2,_ 0.2 mM Na_2_PO_4_, 12 mM NaHCO^3^, 5.5 mM ᴅ-glucose). Neurons were imaged live on a confocal microscope using the 63X objective and 0.66 μs dwell time (488 and 543 nm lasers) with 1X zoom before and 20 minutes after a 3-min exposure to glycine following the same cLTP protocol as described above. Images of dendritic segments with clear spines located proximal to the soma covering a 135 μm x 135 μm field of view were acquired at 0.04-µm intervals in the z-plane.

### Image Analysis and Quantification

Confocal images for a particular experiment were subjectively thresholded using ImageJ (NIH, Bethesda, MA) software and the same threshold was used for all images obtained for a single experiment, throughout the experimental analysis. For live imaging experiments, the same threshold was applied to the images acquired at the indicated time points. Mean intensities over time were then determined using ImageJ. All Z-stack images were Airyscan processed and collapsed as maximum intensity projection using the Zen Black 2.3 SP1 software.

#### Dendritic spines density

Spine density was quantified using the total dendritic length and the total number of spines for transfected neurons. Photoshop CC 2019 software (Adobe Systems Inc., San Jose, CA) was used to create “masks” of the dendrites from the 63X mCherry cell fill. The axon was removed, as well as any dendrites or axons that did not originate from the cell of interest. For all images, the neuron mask was then individually and subjectively thresholded at 180 to binarize the image. Total dendritic length and the number of dendritic spines were analyzed using the default values and analysis for the NeuronStudio (version 0.9.92, CNIC, Mount Sinai School of Medicine, New York, NY) software, adapted from Rodriguez et al(Rodriguez, Ehlenberger, Hof, & Wearne, 2006). Dendritic spines were defined as any protrusion between 0.2 and 3 μm in length emanating from the dendritic shaft, excluding those found on the cell body.

#### SEP-GluA1 insertion

The mean fluorescence intensity of SEP-GluA1 puncta was determined in ImageJ. To quantify the SEP-GluA1 signal, images were thresholded and ROIs were manually drawn around visually identified spines along a dendrite of a transfected cell within the proximal 100 μm from the cell body before and after 3-min cLTP stimulation at the indicated time points. Confocal images shown in the figures were subjected to a 1-pixel Gaussian blur. Levels of brightness and contrast were moderately adjusted in Photoshop CC 2019 software (Adobe Systems Inc., San Jose, CA) using scientifically accepted procedures (Rossner & Yamada, 2004).

#### Colocalization

eGFP-transfected neurons were immunostained for PSD-95 (as a marker of the postsynaptic density) and PRG-1. To only assess the colocalization of proteins within the transfected neurons, a ‘mask’ of the dendrites was created from the GFP cell-fill, imaged at 63X magnification, in ImageJ (NIH, Bethesda, MA). The soma and axon of the imaged cell was removed, as well as any neurites that did not originate from the cell of interest. This cell-fill GFP mask was applied onto PSD-95 positive and/or PRG-1 positive channels to only examine puncta within transfected cells while excluding untransfected neurons. The masks from these channels were then manually thresholded to create binary images. Puncta were defined as being between 0.05 μm and 3.0 μm in size, and puncta density was calculated by obtaining the total number of puncta in the dendrite mask divided by the total length of the dendrite mask. Colocalized points from separate channels were determined using a custom macro incorporating the “Analyze Particles” function of ImageJ and the “Colocalization plugin (https://imagej.nih.gov/ij/plugins/colocalization.html), and the total dendrite length was measured using the NeuronStudio software (version 0.9.92, CNIC, Mount Sinai School of Medicine, New York, NY), as above. Points of colocalization were defined as regions with >50 intensity ratio between the two channels.

### Statistical Methods

Prior to analysis, normality and homogeneity of variance were tested for. All values in the text and graphs are expressed as the mean ± standard error of the mean. *P*-values at or below 0.05 are considered statistically significant. All major statistical analyses, outlier analyses and graph generation were conducted using GraphPad Prism v7 software (GraphPad, CA, United States). Asterisks and hashtags are used to denote levels of statistical significance within all graphs (^∗^*p* < 0.05; \*\**p*<0.01; \*\*\**p*<0.001; #*p* < 0.05; ##*p*<0.01; ###*p*<0.001).

---

**Table S1: LFQ intensities for the 1940 different gene products identified from the LC-MS/MS analyses of (+)HAM and (-HAM) samples from hippocampal membrane fractions of control and fear conditioned mice.**

**Table S2: List of 345 proteins with at least 2 LFQ values in control and FC (+)HAM and (-)HAM) groups**. Fold change in LFQ intensities was calculated between fear-conditioned and control +HAM groups. Proteins are grouped below according to whether they showed increased (pink) or decreased (blue) palmitoylation in the FC group compared to control. Proteins within each group are listed in order by decreasing fold change.

**Table S3**: **List of 121 differentially palmitoylated proteins including proteins that are significantly different between the 2 groups (p< 0.05) and proteins with a fold enrichment ≥ 2 in bold.** Fold change of a protein is calculated by dividing the mean LFQ intensity following FC by the mean LFQ intensity for the control samples. Increased fold change in palmitoylation after FC is shown in pink and decreased fold change is shown in blue. Proteins are annotated by their parent gene ontology (GO) Molecular Function and Biological Function domains. Sub-class represent ’child’ terms that were used for grouping annotations under the parent GO term.

**Table S4: KEGG pathway analysis**. KEGG pathway analysis was performed using g:Profiler, the mouse reference genome, a user significance threshold of 0.05 and the g:SCS method for multiple comparisons correction. Inputs were not ordered, and the domain scope included the only annotated genes.

**Table S5: DisGeNET disease enrichment analysis of the 121 differentially palmitoylated genes**. Terms are ranked in increasing order of *p*-value.

**Table S6: Spearman correlation coefficients for co-expression of 121 substrates (columns) with 30 enzymes/cofactors across 14 cell classes in the mouse hippocampus.** Expression data was obtained from DropViz. Column AG indicates clusters of substrates based on co-expression, while row 125 indicates clusters of enzymes.

**Table S7: Cell-type expression level z-scores for the set of 121 differentially palmitoylated substrates and 30 enzymes/cofactors in the mouse hippocampus**. The results of hierarchical clustering analysis are shown in column O.

**Figure S1:**
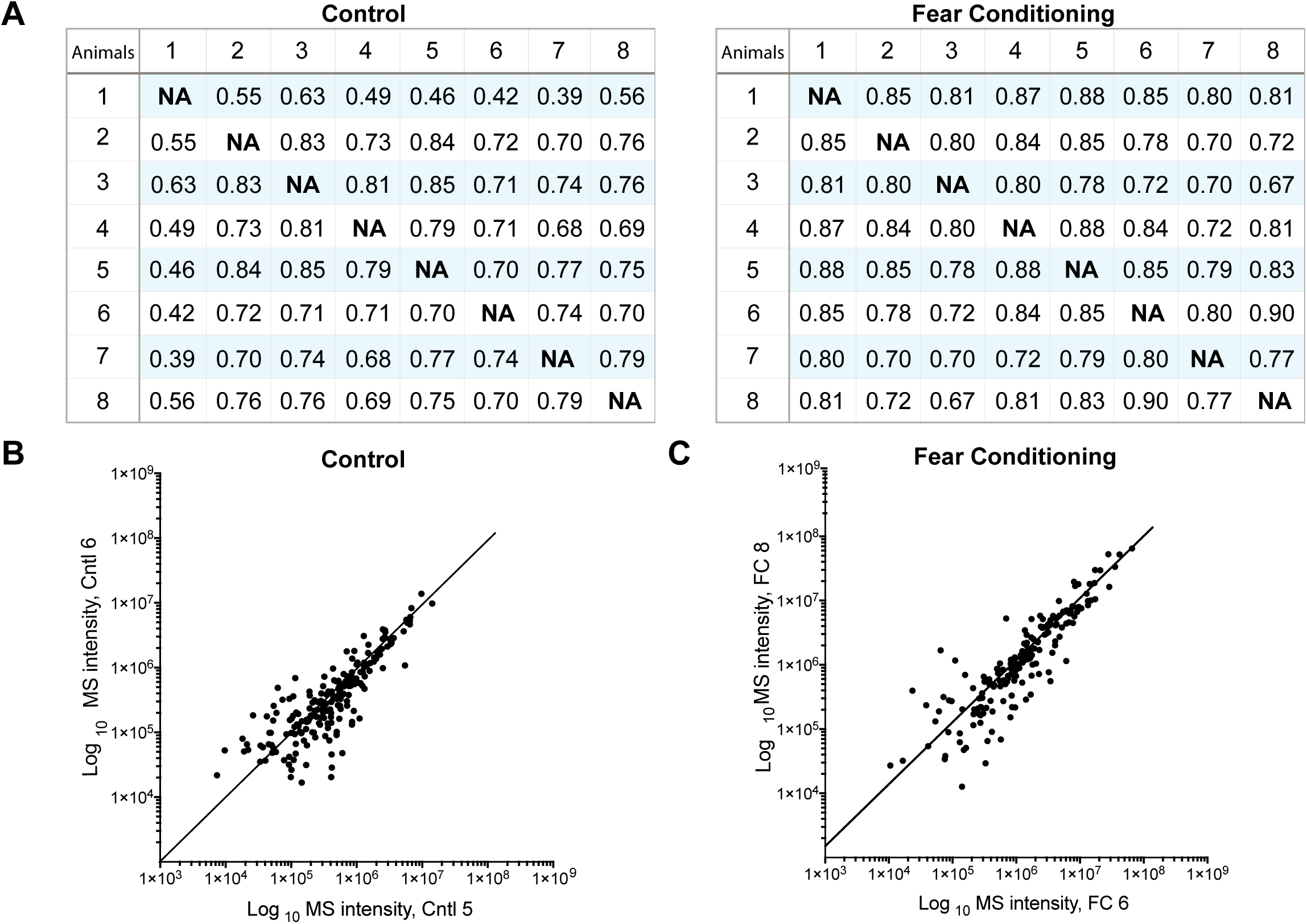
Comparative analysis of sample reproducibility in biological replicates of membrane fractions. (A) Pairwise correlation values between all control replicates and all FC replicates. High r2 values for each comparison demonstrate consistency amongst replicates. (B) Representative correlation of 2 control replicates (Pearson R = 0.70). (C) Representative correlation of 2 FC replicates (Pearson R = 0.90). Correlation of replicates was assessed with Pearson correlation of log-transformed LFQ values. Each dot corresponds to the intensity of the same peptide that was identified in the first (vertical axis) and second (horizontal axis) biological replicates.

**Figure S2:**
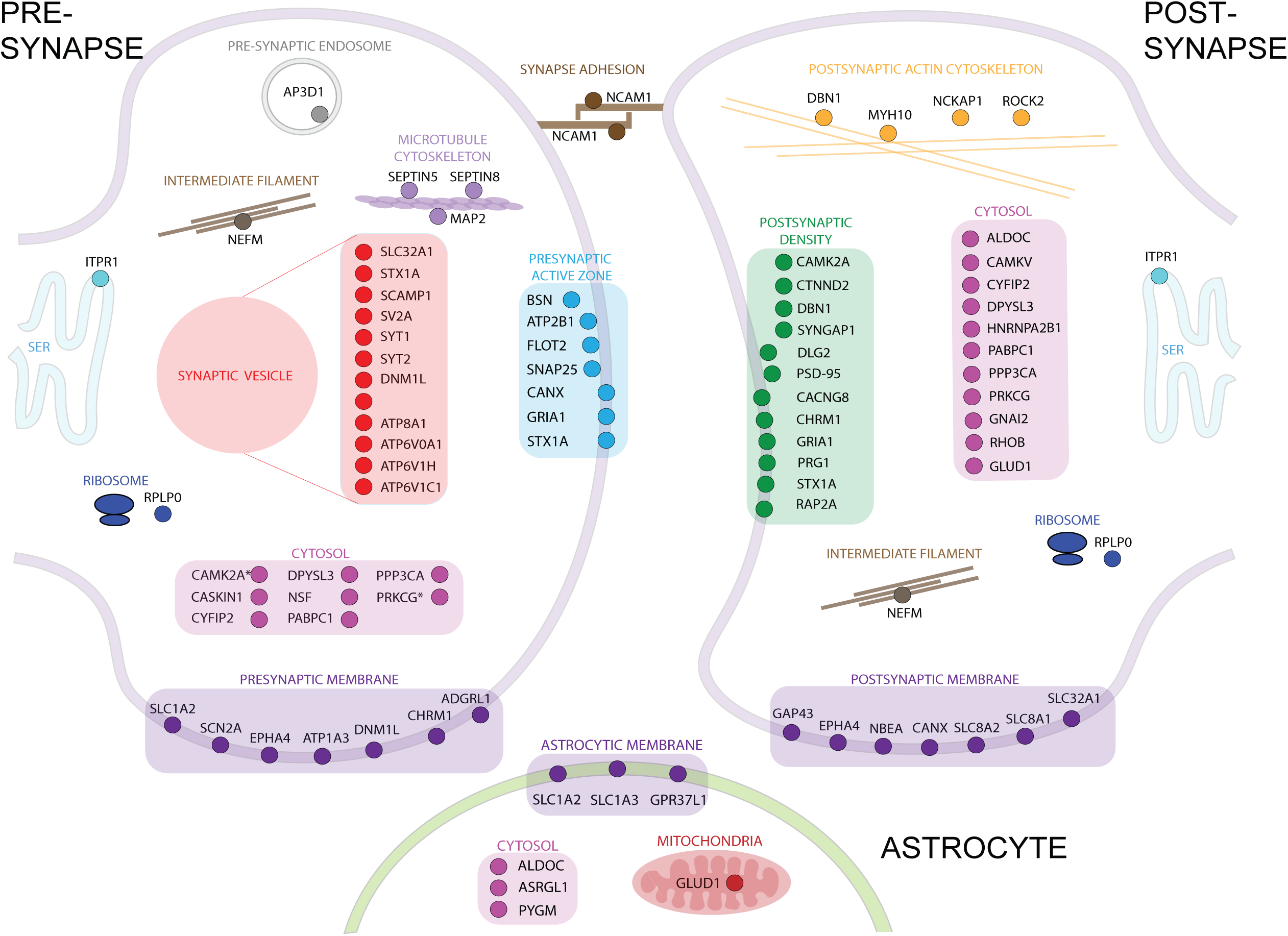
Cellular localization of differentially palmitoylated proteins of interest. Pre- and post-synaptic localization of proteins were made using SynGO (63 proteins) and MGI (70 proteins). 7 proteins were found to be enriched in astrocytes (Astrocyte Transcriptome) in the tripartite synapse.

**Figure S3.**
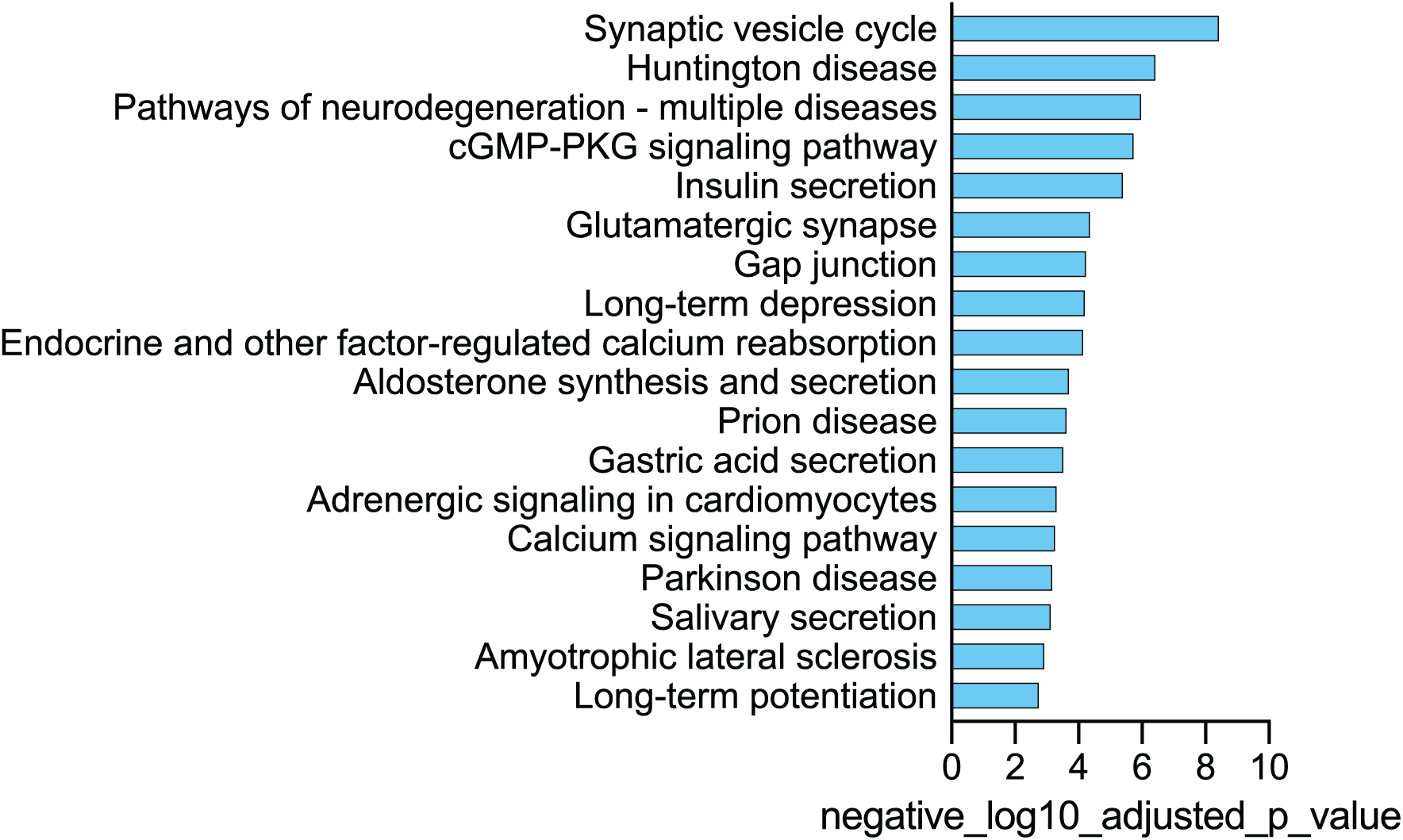
KEGG Pathway Enrichment analysis reveals high levels of enrichment in terms relating to the synapse and neurodegenerative diseases. KEGG pathway enrichment analysis was performed using the web-based program, g:Profiler, with a p-value threshold of 0.05. The 121 differentially palmitoylated proteins were queried within the Mus musculus database. Enriched pathways are shown ordered by the negative logarithm of their p-value.

**Figure S4.**
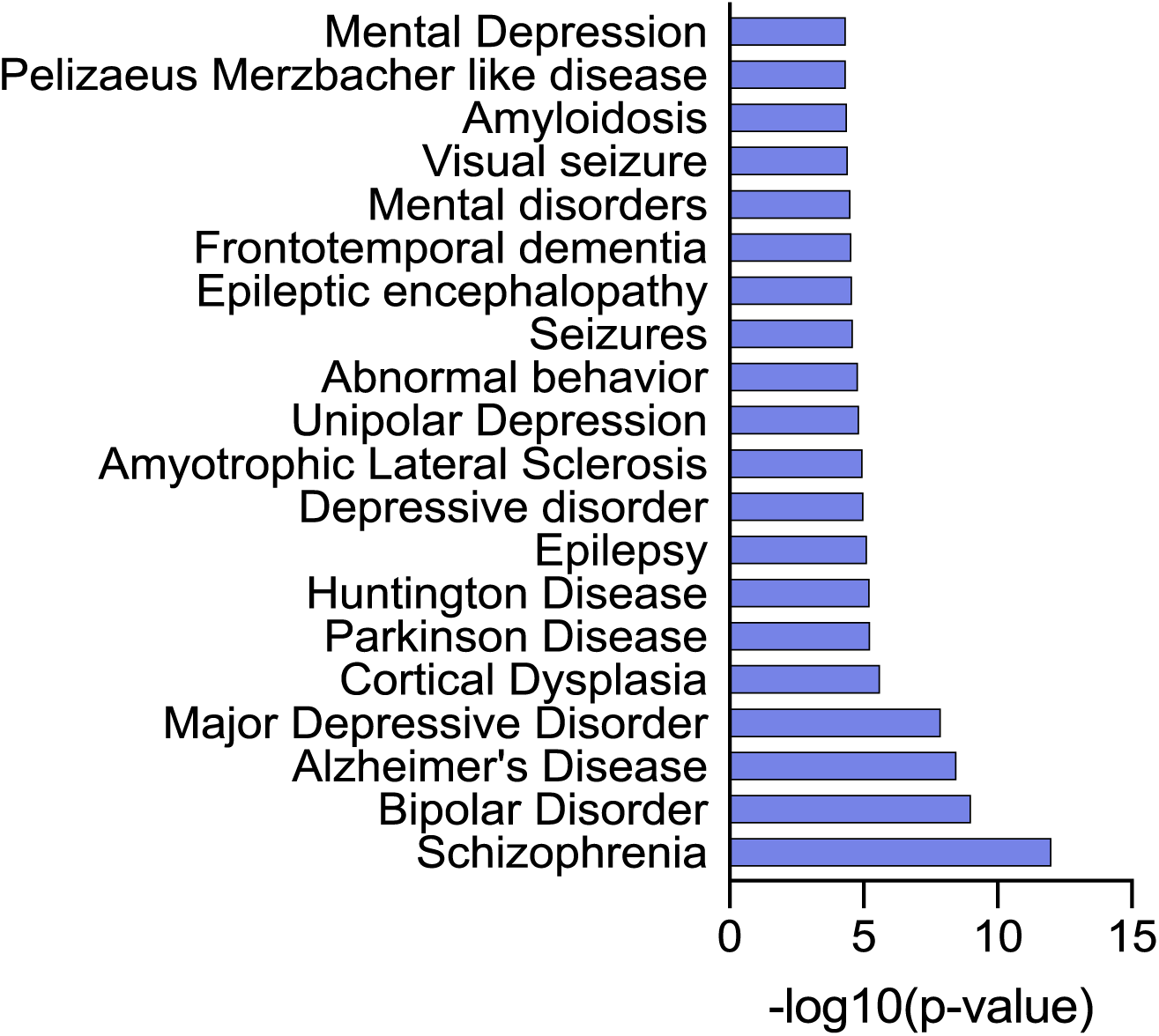
Disease Enrichment analysis reveals high levels of enrichment in neurological and psychiatric diseases. Disease enrichment analysis was performed using the web-based platform, EnrichR and the DisGeNET database. Enriched pathways are shown ordered by the negative logarithm of their p-value.

